# Four classic “de novo” genes all have plausible homologs and likely evolved from retro-duplicated or pseudogenic sequences

**DOI:** 10.1101/2023.05.28.542624

**Authors:** Joseph Hannon

## Abstract

Despite being previously regarded as extremely unlikely, the idea that entirely novel protein-coding genes can emerge from non-coding sequences has gradually become accepted over the past two decades. Examples of “de novo origination”, resulting in lineage-specific “orphan” genes, lacking orthologs, are now produced every year. However, many are likely cases of duplicates that are difficult to recognize. Here, I re-examine the claims and show that four very well-known examples of genes alleged to have emerged de novo “from scratch” - namely *FLJ33706* in humans, *Goddard* in fruit flies, *BSC4* in baker’s yeast and *AFGP2* in codfish - all have plausible evolutionary ancestors in pre-existing genes. In the case of the first two, highly diverged retrogenes that code for regulatory proteins may have been misidentified as being orphans. The antifreeze glycoproteins in cod, moreover, are shown to have likely not evolved from repetitive non-genic sequences but, as in other related cases, from an apolipoprotein that may well have been pseudogenized before later being reactivated. These findings detract from various claims made about de novo gene birth and show there has been a tendency not to invest the necessary effort in searching for homologs outside of a very limited syntenic or phylostratigraphic methodology. An approach used here for improving homology detection draws upon similarities, not just in terms of statistical sequence analysis, but also with biochemistry and function, in order to obviate failure.

## Introduction

The discovery of many lineage-specific genes across eukaryotic genomes has been a remarkable development (Wissler et al 2013). Although some can readily be explained in terms of duplication and divergence (Johnson 2018; Weisman et al 2021), others have apparently arisen “de novo” from DNA sequences previously not coding for proteins (Wilson et al. 2017; Van Oss and Carvunis 2019). In such a scenario, new genes can be created entirely “from scratch” (Schlötterer 2015; Vakirlis et al. 2018). Genomes may be potentially teeming with emerging “protogenes” (Carvunis et al. 2012; Grandchamp et al. 2022), thus representing an important part of evolutionary innovation as well as adaptation (Guerzoni and Mclysaght 2015; Blevins et al. 2021).

The basic idea behind “de novo origination” is that intergenic sequences, that are known mostly to be transcribed, albeit at lower levels than coding regions (Hanguauer et al. 2013), will contain open reading frames (ORFs). Most of these are expected to be quite short, but one or more mutations could extend them to the point that they would be comparable in size with most genes. If they are translated into proteins, they could be preserved by selection should they be both functional (including folding properly) and biologically useful to the organism. Even if they do not confer any function, but do not interfere with other proteins, they could still become fixed due to drift. Until relatively recently, this was thought to be next to impossible. However, as sequencing and comparative analysis progressed, researchers found evidence there were far more lineage-specific genes than was expected and reasoned that some may be explained by stretches of non-coding junk DNA becoming translated (Tautz 2014).

The identification of candidate de novo protein-coding genes is complicated by the presence of genome assembly errors (Zhang and Backström 2014), which are surprisingly common, and can lead to the false interpretation that a gene in other lineages is not translated. There is often not sufficient proteomic data to confirm that a protein is only being synthesized in one lineage to the exclusion of others, even if the observed sequence data appears to preclude such a possibility. Two protein coding genes previously identified as having arisen de novo in the human lineage (Knowles and Mclysaght 2009), namely CLLU1 and C22orf45, now lack peptide status according to PeptideAtlas (Desiere et al. 2006; Wu et al 2011) and are likely lncRNAs. Even if their status were to change yet again, only the verified absence of the gene products in chimpanzee tissue would support the claim that they are human-specific.

Furthermore, many long ncRNAs have been found to be translated, whilst some transcripts of protein-coding genes are not (Hartford and Lal 2020), obscuring the distinction between coding and non-coding regions. These long ncRNAs are now proposed as being a potential intermediate stage in the origination of a de novo protein-coding gene (Ruiz-Orera et al. 2014; Tautz and Domazet-Loso 2011). Some may, however, be formerly coding pseudogenes that have assumed a new role as regulators of gene expression (Pink et al 2011; Milligan and Lipovich 2015). Another alternative to a hypothesis of de novo birth is the parallel deactivation of genes in multiple lineages except for just one (Siepel 2009). This would appear to be unlikely, especially if the same disabling mutations are present in more than two lineages, although mutational hotspots/biases (Svensson and Berger 2019) can result in the homoplastic divergence from a functional precursor, meaning this is a real possibility.

The resurrection of pseudogenes (Brosch et al. 2011; Prade et al 2018) is yet another intriguing scenario whereby a gene long dead has been reactivated by enabling mutations. Indeed, the non-coding DNA that de novo genes are said to emerge from is littered with sequences that were formerly coding (Zhang et al. 2003). These include processed pseudogenes that are reverse transcribed and randomly inserted into the genome, lacking both introns and promoters (Ciomborowska et al. 2013). It is no small coincidence that many de novo genes also lack introns and are often dependent on the promoters of other genes for transcriptional activity (Gotea et al. 2013). In a study reporting candidate de novo genes in Oryza, Zhang et al. (2019) admit they had to exclude cases of functionalized former pseudogenes liable to be mistaken for de novo genes. Casola (2018), moreover, found that hundreds of de novo genes identified by phylostratigraphy may, in fact, really lie within the sequences of old pre-existing genes.

Because gene duplicates move about the genome through ectopic recombination and retrotransposition (including exon shuffling), a syntenic analysis that Vakirlis et al. (2020) recommend is unlikely to prove that helpful in identifying sequences that have not previously coded for proteins. Pseudogenes are expected to diverge substantially given the respite in selective pressure, but if they have not completely degenerated by mutation, and manage to retain something of the original gene function, they could still prove useful to the organism if ever brought back to life (Troskie et al. 2021). When this happens, as unlikely as it is, any BLAST search may not pick up on this and so they will appear to have emerged “from scratch” even though this would be a case of homolog detection failure, as explained by Weisman et al. (2021). Marsch-Martínez et al (2022) admit when reporting a possible de novo gene within *Brassicaceae*, that “the identification of a lineage-specific gene that has originated de novo could be the result of a homology detection failure.” In a “conservative” review, covering just a handful of de novo protein-coding genes whose peptide status and function has been demonstrably established, Weisman (2022) also acknowledges that, for some of the reasons as listed above, there is likely to be a significant overestimate of the number of candidate de novo genes. The phenomenon of stop codon read-through (Li et al 2019), as well as alternative start sites (Bayzkin and Kochetov 2011; Rojas-Duran and Gilbert 2012) also means that sequences that did not look like having viable ORFs in some lineages may, in fact, be translated in their entirety by the ribosome in the same way as those with uninterrupted reading frames. Parallel gene loss across multiple independent lineages is, however, still considered to be much less likely an event compared with gene gains arising from previously non-genic DNA in a single lineage.

This article focuses on a selection of extensively cited examples of “de novo origination”, two of which Weisman (2022) draws specific attention to. It attempts to show that, in each case, plausible alternative accounts exist that have either not been identified or unduly dismissed as not being parsimonious. Other candidate de novo genes were considered for inclusion, but the ones examined here taken from four species (human, fruit fly, yeast and cod) were found to be of most scientific interest. They could, thus, serve as being generic examples, rather than odd exceptions to the rule of limited use to the study of the evolutionary origins of novel genes. A provisional methodology is also proposed to help detect any cryptic homologs and to identify certain instances where a hypothesis of de novo origination is likely to be better explained by other phenomena that would mainly involve duplication and divergence.

## Homology Detection

Homology detection is itself an inexact science that can involve following up on discrete clues (Madej 2007). Deterministic heuristic algorithms like BLAST or HMMER can be fraught with false negatives even if the parameters used in a search are relaxed. The absence of evidence for detectable homologs in outgroups cannot be confidently claimed as evidence that homologs are absent (Weisman et al 2021; Patraquim et al 2022): a gene may have duplicated and diverged rapidly, resulting in a failure to detect orthologs (Broeils et al 2023; Basile et al 2019). A local alignment tool like BLAST finds regions of most similarity and filters results from a public database using a calculated score. However, any natural gaps, owing to indels, and other incongruities, potentially arising from where two related sequences have diverged substantially, can reduce this score considerably (McLean 2004). Penalties introduced for the presence of gaps are important for the sensitivity of the search tool, namely the ability to find any remotely related sequences (Goonesekere and Lee 2004). However, long gaps can also be overpenalized resulting in alignments that may not be biologically correct (Mott 1999).

Accordingly, the E-values, that are often interpreted as statistical differentiators between homology and coincidence, may turn out to be misleading (Ochoa et al 2015). This is especially true where the sequences have diverged because of compensatory changes involving substitutions at sites other than where deleterious changes have occurred (Hartl and Taubes 1996): BLAST uses substitution matrices that score not only by an exact match, but also by the chemical similarity of each aligned amino acid.

One approach, analogous to the use of N-grams, is to focus on just part of the protein sequence (Menlove et al 2009; Atlschul et al 1997). Indeed, particular interest often focuses upon only a small region, domain or pattern within a sequence (Zhang et al 1998). The BLAST algorithm is sensitive enough to pick up on commonalities that would likely be ranked low down or even not returned if the entire sequences were compared. Although this increases the elasticity of any search, it does also have the drawback of potentially producing spurious hits that may not indicate any direct ancestral relationship. This is often the case with iterative approaches like PSI-BLAST (Altschul et al 1997) or DELTA-BLAST (Boratyn et al 2012) used in remote homology detection where false positives can abound when very short sequences are involved. Such hits need not be due necessarily to coincidence, but to very indirect relationships produced by more ancient duplications or instances of illegitimate recombination. Moreover, only certain combinations of amino acids confer function because of various biochemical/biophysical rules. Therefore, discretion needs to be exercised when searching for any homologs where there may well be a high level of divergence.

Lee et al (2008) explain that in the mathematics of BLAST, the E-value (*E*_seq_) for a single sequence comparison, i.e. not including all subject sequences contained in the public database, can be calculated as shown below. This value is reflective of the expected number of local alignments with a bit score of at least S. Here, L is the length of the query sequence aligned against a subject sequence that is of length M. The parameters K and λ are natural scales for the search space size and the scoring system respectively.

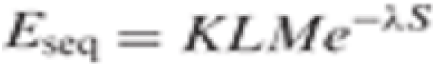

This expectation value can be informative when evaluating the similarity of any two sequences where reference to all other potentially similar sequences is irrelevant or unhelpful. But where the public database, containing N amino acids within the protein repertoire, is included in the calculation of the E-value (*E*_db_), then M is replaced by N. The E-value below is thus a corrected bit-score adjusted to the sequence database size:

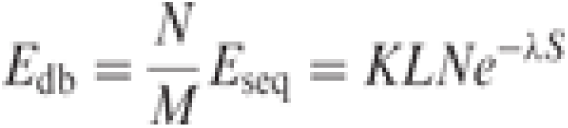

The effect of reducing the query length, namely by only taking a subsequence from it, serves to also reduce the effective number of independent tests performed by BLAST. In some cases, a potentially more realistic E-value can, thus, be determined by a factor inversely proportional to the original query length. This is very useful where the query subsequence is similar to most of the subject sequences, but the rest of it is not. If the entire query length were used, then the resultant E-value could be much greater. As such, in addition to relaxing parameters to conduct a whole sequence query search, focusing just part of the query sequence, either of the most interest or likely to more conserved, can be useful. The N-terminus is usually a good choice since this region usually, but not always, contains the important signal peptide for those proteins that are secreted or translocated to an organelle within the cell (Choo et al 2009). The demarcation of the N-terminus varies considerably but, for the purpose of a sub sequence search, a length of *L*/6 amino acids from the start site is a fair approximation for proteins that are greater than 100 amino acids. It is also to be expected that any two homologs should, in principle, be align-able at their respective N-termini unless one of them has experienced a bulk deletion therein. Whilst frameshifts do naturally occur at the N-terminus, they are likely to be greatly more deleterious than those downstream at the C-terminus where any stark sequence divergence and premature termination is tolerated much better, especially for duplicates (Okamura et al 2006). The N-terminus is thus found to be generally, but certainly not exclusively, more conserved than the C-terminus (Schejter and Shilo 1985; Phansalkar et al 2013; Alawad et al 2016). A good example is the case of antifreeze glycoproteins in Antarctic notothenioids that appear homologous with trypsin and encoded by genes proximate to each other. Very close similarities were noticed at the N-terminus, specifically with the signal peptide, even though the remainder of the sequences downstream are markedly different (Chen et al 1997). As a result of sharing this specific feature, both gene products are secreted within the pancreas. Such an approach was used to identify a relationship between histones and ribosomal proteins (Bozorgmehr 2019).

As well as the E-values, homology may be adjudged using the overall level of sequence identity with a global alignment which is not assessed using a local alignment tool. Generally, any two sequences sharing an identity >30% most probably have a common origin, although this can be incorrect when they are short or if their composition is similar (Pearson 2013). E-values, of course, can still be low when there is still a relatively low level of identity, as in a case of a region that has been conserved in otherwise divergent sequences, and high where the identity is also high. The size of the database, and bias for certain combinations, can be yet another important factor. Moreover, some proteins can share a sequence identity as low as 15% and still be recognized as homologous in terms of their structural congruence as reflected in the Root Mean Square Deviation (RMSD), a measure of the similarity of superimposed atomic coordinates, where a value of < 3 angstroms is regarded as a close match (Reva et al 1998). However, a disadvantage of the RMSD is that it is dominated by the largest error and so is strongly affected by the most deviated of the fragments and by protein size – models have been designed to attenuate this shortcoming (Kufareva and Abagyan 2015). Without biological insight into what a gene might be related to, conducting an exhaustive database search in terms of structure, with pre-computed sequence-structure or geometric alignments, can prove difficult. A useful algorithm is VAST (Madej et al 2014), which identifies homologs that cannot be recognized in a sequence comparison by availing of a molecular modelling database. Another criterion is the likelihood that any two sequences are related because of shared functional roles they have (Barthez et al 2020; Struhl 1987 that tend not to change significantly. Also important is the exon-intron organization, location on the chromosome and where peptides are synthesized and localized to in the cell. Multiple lines of evidence, including the use of experimental studies, are required to validate such relationships.

Proteins identified as originating de novo may be disputed in terms of their amino acid composition and not just the presence of possible homologs. Previous research has confirmed that natural proteins, unlike random heteropolymers, do not conform to the expected frequency distribution for amino acids based on the genetic code (Bozorgmehr 2015). Although this can be affected by the underlying GC content of the DNA coding for the protein, the extent to which the distribution deviates from the genetic code may serve to detract from an explanation of de novo origination from unstructured and unconserved intergenic DNA. P-values can be calculated using a chi square test to determine whether the observed distribution is statistically significant.

## Results

### FLJ33706: The case of a revived pseudogene?

FLJ33706, also referred to as C20orf203, is a human-specific gene that is highly expressed in the brain and implicated with nicotine addiction according to GWAS studies that identified an SNP in the 3’UTR (Li et al 2010). It is regarded as a *bona fide* de novo human gene that emerged from ncDNA. The orthologous locus in the greater and lesser apes is interrupted with stop sites; this suggests that it is unlikely that the gene product is viable in these lineages, although the possibility of a read-through exists and the absence of a peptide in chimpanzee tissue has not been determined. Transcripts exist in other eutherian mammals meaning that the sequence itself may be at least several tens of millions of years old. The gene has no detectable homologs and an Interpro search could not find any functional motifs shared with other genes.

An alternative possibility is that FLJ33706 is derived from a duplicated pseudogene that has been reactivated in the human lineage alone. As acknowledged, rates of nucleotide substitution in pseudogenes can be quite high due to the respite in purifying selection (Li et al. 1981; Bustamente 2002). A BLAST search would struggle to find any homologs if the sequence has itself diverged significantly during its inactivation. Using adjusted gap penalty parameters and a relaxed Expect threshold, as explained in the Methods section, the entire sequence was then BLASTed against the human proteome. Position-Specific Iterated BLAST was used as it is sensitive to weak but biologically relevant sequence similarities (Altschul et al. 1997): in the first iteration, which is identical to BLASTP, a number of potential hits with E values <1 were returned. Notably, one of them was for a zinc finger protein, the longest isoform of ZNF678, which is interesting given that experiments in rats have shown that these transcription factors are implicated in addiction/aversion to drugs similar to those with the effect of nicotine (Chandrasekar and Dreyer 2010). The region showing the most sequence similarity is at the N-terminus of both genes. In terms of both relative position and sequence coverage, the zinc factor protein appears to be most plausible. Subsequent iterations of PSI-BLAST preserve ZNF678 as being one of the top listed hits.

As explained above, a subsequence search can, perhaps, be more revealing than using the whole sequence. Taking the first 32 residues at the amino end of the 194 aa protein, corresponding to a sixth of the overall sequence, this was BLASTed against the human proteome. A few interesting matches above the default threshold were produced for several functionally diverse genes. This is to be expected since, according to the general prediction of gene duplication, most genes should share characters in common with each other either due to a shared phylogeny or lateral acquisition (e.g. exon shuffling). The zinc factor N-terminus produced an E value of 0.0006, easily below the default threshold of 0.005; in terms of overall sequence identity, FLJ33706 can be aligned in its entirety with the 580 aa zinc finger protein to give an overall identify of exactly 25%, with 61 residues matching. This is displayed in Figure 1. It includes some notable sequence gaps which could be indicative of large deletions in the former. The alignment would fall within the “twilight zone” that denotes a range (20-30% identity) where the signal is such that homology *could* exist (Khor et al. 2015; Pearson 2013). A 25% cut-off was used for orthologs considered present in the LUCA (Weiss et al 2016).

**Figure 1:**
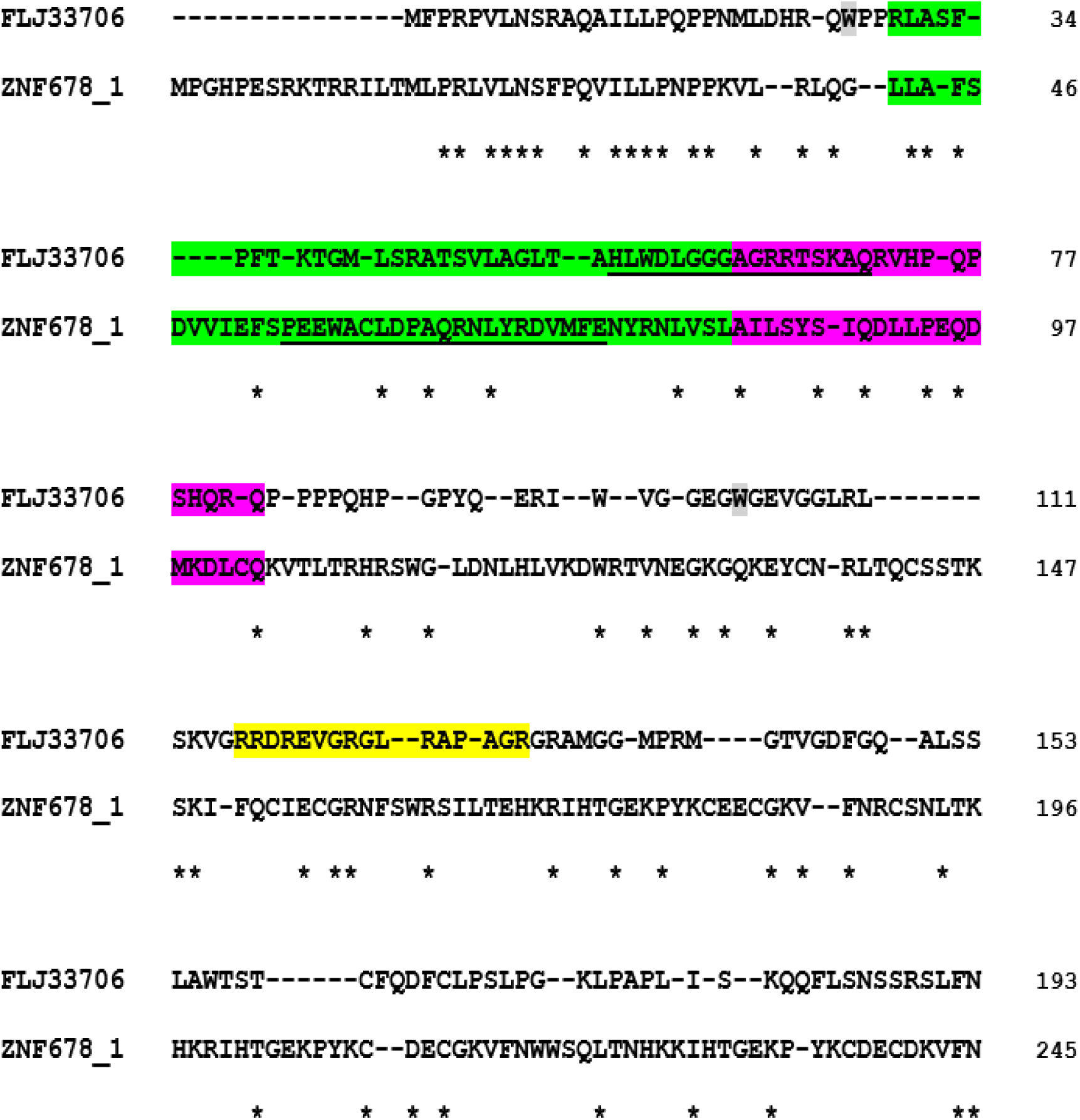
The alignment of FLJ33706 and ZNF678 (isoform 1) in humans. Asterisks denote the shared residues. Overall, there is a 25% identity inclusive of some significant gaps. The zinc finger protein is 580 aa long, and the rest of its sequence is not shown. Shaded in green is KRAB A-box, and in pink is KRAB B-box, in ZNF678.1 along with the corresponding sequence in FLJ33706. Underlined are the residues most structurally aligned. Shaded in grey is a residue that was originally a stop site at the N-terminus but was replaced with tryptophan in the human lineage alone. Another one further downstream is where an insertion was also deleted, again only in humans, allowing the reading frame to avoid other stop sites and so to continue. Shaded in yellow is a 16 aa arginine-rich localization signal that is predicted for nuclear import.

If, indeed, ZNF678 is the precursor of FLJ33706, then a premature truncation or bulk deletion at the C-terminus would easily explain the much shorter length. The N-termini, however, are not fully aligned, with the first 15 residues of the zinc finger protein apparently absent in FLJ33706. As displayed in Suppl Fig 1, for the first 22 codons downstream of the methionine start site in FLJ33706, 11 of them (i.e. half) are identical with those that are aligned with the corresponding sequence in the zinc finger protein. Another 3 are synonymous, whilst the remaining 8 code for different amino acids; of these, 5 differ by just a single nucleotide substitution. Codon usage analysis reveals there is an over-representation of the frequency of the CCT codon (proline) in the N-termini of both genes than would be expected (Parvathy et al. 2022).

The 90 nt sequences (including gaps), shown in Suppl Fig 1, are 68% identical which produces an E-value of 1 * e^-14^ for the pairwise alignment using BLASTN, albeit independent of the public database. As explained, the initial 15 codons situated at the N-terminus of ZNF678.1 are missing in FLJ33706, which resulted in a new start site further downstream. One possibility to account for this observation is that they were simply lost due to a bulk deletion. Another is that they might have been pushed into the 5’UTR, and so do not code for amino acids any longer. By inspecting the nucleotide sequence of the 5’UTR of FLJ33706, it is evident that the latter has likely happened, and it is even possible to reconstruct the events concerned. Suppl Fig 2 shows the alignment of the amino end of ZNF678.1 against the terminal 5’ UTR nucleotide sequence of FLJ33706. The aligned sequences are as much as 57% identical and include at least 7 indels. BLASTN, however, still does not find this to be significant when a pairwise alignment is conducted, despite it being a higher level of identity than as would be expected for two unrelated nucleotide sequences (ranging from 10-50%).

However, there is another indication, beyond a purely statistical approach, that the 5’UTR of FLJ33706 contains the remnant of the amino end of ZNF678.1. As can be seen, guanine appears to have been inserted in the methionine start codon of the latter when compared to the corresponding sequence of the former. This would have had the effect of creating a frameshift that pushed the start site downstream in the case of the putative daughter gene. In the zinc finger protein, the CCA (proline), and part of the GGA (glycine) codons, subsequent to the start site, are evident in the 5’UTR of FLJ33706. In addition, it appears that a Kozak sequence *(GTCTCCTATG*) was formed through two deletions. Li et al. (2010) posit that FLJ33706 is a gene of viable length partly because of a substitution (T**A**G->T**G**G) close to the N-terminus that resulted in an in-frame stop site being replaced with the codon for tryptophan. In the other great ape species, the stop site still exists (assuming there are no assembly errors) which means that, even if transcribed, the resulting gene product would be too short to prove functional. In the zinc finger protein, the corresponding codon is GGA (glycine).

They also note that there was also a 1 bp deletion (guanine) 104 codons downstream of the start site, which also may have caused the reading frame to then escape other potential stop sites, and postulate that the resultant frameshift in the human lineage produced a unique encoded amino acid sequence. However, a comparison of both genes reveals that the corresponding sequences still show some similarity close to the C-terminus of FLJ33706. Therefore, it is more likely that the deletion is itself a back mutation in humans which served to restore the remainder of the ancestral reading frame, although still largely diverged, from a prior insertion present in other primates whose nucleotide sequences are otherwise very similar to the one in humans. There is evidence of reading-frame-restoring mutations in other species (Esfeld et al 2018).

Organizationally, FLJ33706 is unusual for a candidate de novo gene in that it contains four to six exons depending on which of its respective splice variants is observed. Li et al. (2010) claim that the repeated insertion of *Alu* sequences accounts for these intronic sequences, as well as part of the coding sequence also. Both the 5’UTR and 3’UTR sequences are interrupted by the presence of two large introns within them. The coding sequence in FLJ33706 is itself separated by an intron which is rather short in length. The zinc finger protein contains three introns in total, including one just downstream of the translational start site corresponding to one of two in FLJ33706 that interrupt the 5’UTR. As stated above, this contains the remnant of the N-terminus of the zinc finger protein. The splice sites and position of the intron in the respective sequences are also shown in Suppl Fig 2. Despite having similar length and structure, there is nonetheless little in common between the actual nucleotide sequences for the corresponding introns. Therefore, an inference of *Alu* element insertions is quite likely given the differences both in terms of nucleotide arrangements and in intron length. If FLJ33706 is indeed a retroposed copy, it would likely have had no introns initially, having been spliced out prior to reverse transcription, and so would have had to have acquired them. Intron gain has been inferred in the case of some retrogenes (Kang et al. 2012). Preferential insertion points (Comas et al. 2001) could explain why *Alu*-derived introns appear in their respective positions in FLJ33706 such that they resemble those for ZNF678.1. The apparent insertions are also contained in other apes.

The primary sequence of FLJ33706 would appear to correspond to that of the Krüppel associated box (KRAB box) domain at the amino end of the zinc factor. This is used for transcriptional repression by many zinc finger proteins, whereby the domain functions by binding with co-repressors through protein-protein interactions (Margolin et al 1994). KRAB domains are typically 75 aa in size and contain a KRAB-A box (40-50 amino acids) that is both necessary and sufficient for transcriptional repression, along with a KRAB-B box (20-25 amino acids) having an accessory role. The KRAB boxes have been largely preserved in FLJ33706, and with key leucine, glutamine and phenylalanine residues still present within them. FLJ33706, however, differs markedly from the corresponding peptide sequence of ZNF678.1 in terms of its amino acid composition in that it is distinctly more alkaline. There is also an abundance of glycine and proline residues that are often found in turn and/or loop structures (Krieger et al. 2005). This is reflective of the relatively high GC content of the gene, compared to its putative ancestor, which could indicate a large degree of GC-biased recombination over time.

A comparative sequence analysis can take us only so far when looking for homology. Any two proteins can be highly divergent in sequence, but similar in terms of their three-dimensional structure, reflecting not just shared function but a common origin also. Figure 2 shows the predicted structure of the N-termini of FLJ33706 and the zinc finger. As with all KRAB box domains, it consists of up to two amphipathic alpha helices, with the remaining structure being loop-like. A structural alignment, focusing just on the KRAB domains, produces a an RMSD of 2.93 Å with 17 residues, shown in Figure 1, appearing to be critical to conformations displayed in Suppl Fig 3. As a benchmark, the RMSD of the KRAB domain in the zinc finger and that of its consensus sequence was determined to be 2.15 Å. The residues that are predicted to interact with DNA for both FLJ33706 and the corresponding sequence of ZNF678.1 are displayed in Suppl Tables 1-2, with functional predictions shown in Suppl Tables 3-4: FLJ33706 is also computationally predicted, with more probability, to be a transcription factor than ZNF678.1, and with a greater proportion of its residues binding with DNA. Deep learning classification models also corroborate this finding. A predicted downstream nuclear localization signal is shown in Figure 1. This would allow the protein to enter the nucleus and bind with DNA. However, despite these features, FLJ33706 was still determined to be up to 40% disordered compared with the corresponding sequence of the zinc factor that is just 10%. One claim that is often made about de novo genes is that they are not sufficiently ordered to be subject to the same level of aggregation as most other proteins (Wilson et al 2017), and this might permit structural flexibility.

**Figure 2:**
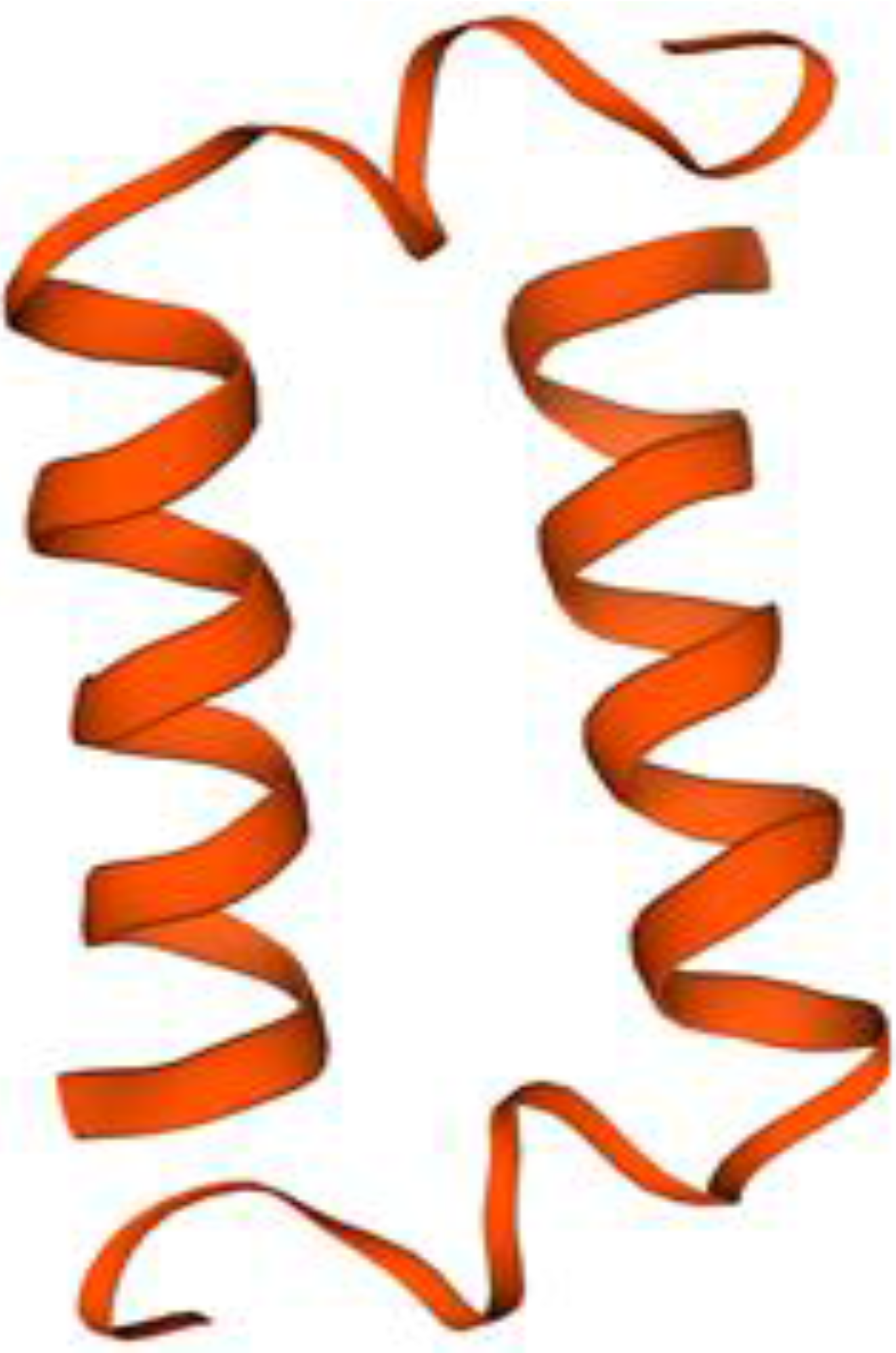
The computationally predicted structure of the amino end region of FLJ33706 (corresponding the KRAB domain sequence in ZNF678.1) revealing a typical Krüppel box conformation consisting of exactly two alpha helices with the rest of it being disordered loop.

**Figure 2b:**
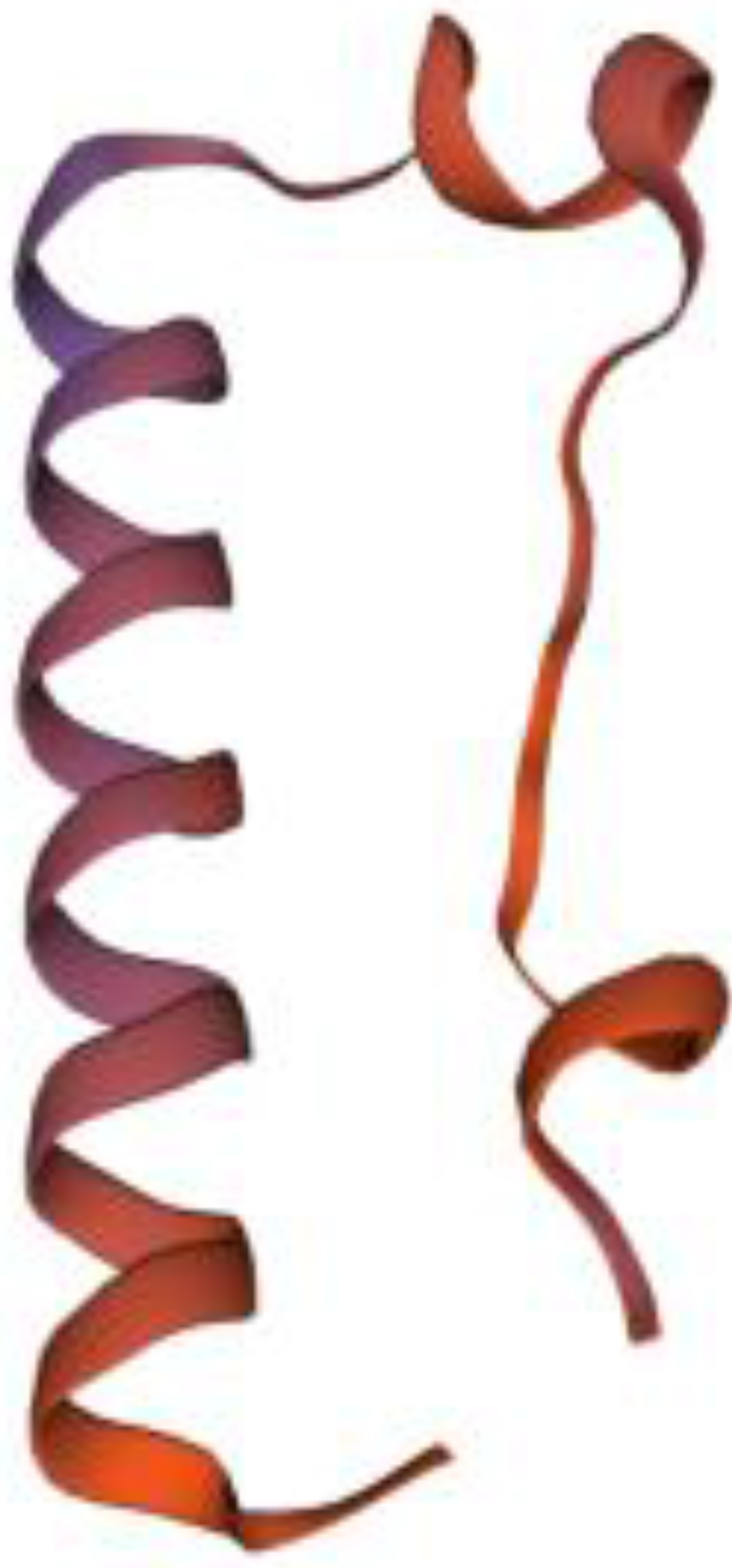
The predicted structure of the part of ZNF678.1 for the N-terminal KRAB domain.

The expression patterns of both genes reveal both similarities and differences. FLJ33706 is highly expressed in the brain whereas the zinc finger protein is found there, including the cerebral cortex, but is also expressed within the testes, prostrate and thyroid gland (Uhlén et al. 2015). Li et al. (2010) speculate that FLJ33706 depends for its transcription on genes flanking it on the reverse strand, along with the open chromatin structure. They are situated on different chromosomes, namely 1 and 20, which indicates that the latter would be the result of retrotransposition. Indeed, retrogenes exhibit a higher incidence of tissue-specific gene expression than their parents do (Carelli et al. 2016). The zinc finger protein likely has more distal enhancer interactions. There is evidence that the KRAB domain in zinc finger proteins plays a key role in neuronal development (Farmiloe et al. 2020; Turrelli et al. 2020). The KRAB ZNF gene family is also known for its rapid evolution in hominids (Nowick et al 2010).

In conclusion, it is perfectly plausible that FLJ33706 is a retroposed and truncated copy of one of the isoforms of a zinc finger transcription factor. Through several changes, an initially inactivated retrogene eventually became an active human-specific protein coding gene. This type of event is not well-attested, and most likely rare, but still theoretically plausible. This makes much more sense than the alternative scenario that involves the recruitment of an unspecific stretch of intergenic DNA – of undetermined origin and function (if it had any) that may have become a lncRNA, as an intermediate stage, before being translated as a protein. The extent to which the gene contributes to human cognition is unclear, although other “brain paralogs” are present in *H. sapiens* (Fiddes et al. 2018; Florio et al. 2015) and are considered to have some effect.

### GODDARD: a “putative” de novo gene?

Another candidate de novo gene, Goddard, is found exclusively in the Drosophila genus, but not among all its species. Its de novo status is referred to as “putative” because high levels of divergence, “prevent the unambiguous identification of the (related) non-coding sequences” in outgroup species (Gubala et al 2017). The identification of a non-coding orthologous region in other species is required by McLysaght and Hurst (2016) to demonstrate a definitive de novo origin. Even so, as is consistent with such an origin, Goddard is predicted to encode a protein that has no sequence similarity to any annotated protein. An Interpro and conserved domain search was conducted that also found nothing at all. However, as Gubala et al. (2017) also admit, a duplicated or laterally acquired gene has the potential to evolve rapidly, and so lose any detectable similarity such that it is falsely classified as a de novo gene (Moyers and Zhang 2016). They admit it could still be present outside of Drosophila.

Goddard is reckoned to be about 50 million years old and appears to be required for spermatogenesis where it localizes in the cytoplasm to regulate sperm axoneme assembly (Gubala et al. 2017; Lange et al. 2021). It must have become essential despite its purportedly de novo origin. Interestingly, Goddard is a single exon gene, consisting of 114 codons in D. melanogaster, that exists within a large intron of a well-conserved gene, Omega. Another de novo gene cited along with Goddard, namely Saturn, is not considered here. In this respect, it is similar to PBOV1 in humans (Samusik et al. 2013), a candidate de novo gene also located within an intron. Since Goddard is present in all but one Drosophila species, it makes sense to look for a possible ancestral precursor in another related major dipterid genus. *Musca,* which includes the common house fly, *M. domestica,* is an appropriate taxon to use for the search. As with FLJ33706, the N-terminus of Goddard in the model species, D. melanogaster, consisting this time of the first 20 residues, was BLASTed to see whether the algorithm picked up anything based on this partial sequence. Also, as Gubala et al. (2017) had done, it was searched against the medfly, *Ceratitis capitata.* One of the hits was found to be interesting because the N-termini of both subject and query are aligned in the same position. Although worse than the default threshold, the E-value is still <1. The gene, Protein MAX, is a conserved 165 aa transcription factor which is deployed in several structures and has a clear ortholog in humans. It is one of a number of helix-loop-helix proteins. Subcellular localization prediction, using a machine learning approach unlike the rules based one as deployed by BLAST, shows that the most relevant match to Goddard is actually MAX: the others are localized either to the nucleus or to the cytoplasm. The fact that Goddard is observed at the axoneme is not so unsual; Sdic in D. melanogaster is an axonemal dynein whose parent, Cdic, is a cytoplasmic one (Ponce and Hartl 2006).

If the entire sequences are aligned, inclusive of gaps, they are 30% identical, with 39 residues matching. This is shown in Figure 3. About 9 times out of 10, a 30% identity can be attributed to homology (Rost 1999). One possible exception, stated above, can be when both proteins have a very similar amino acid composition. However, Goddard is a very negatively charged protein, with acidic outnumbering alkaline residues by 2:1, whereas they are finely balanced in MAX. A cluster of aspartic acid residues located at the N-terminus is the most notable feature in common, which was what BLAST picked up on. Moreover, key arginine residues, that are known in other genes to bind both to the negatively charged phosphate backbone in DNA, as well as other amino acids, have been both retained and gained. Glutamic acid and asparagine are also over-abundant with leucine under-represented. The genes exhibit a strong bias for the GAC (D), TTC (F) and AAC (N) codons. A chi square test for the amino acid composition produces a significant P-value < 0.01 meaning that it deviates from the random expectation of the frequency of amino acids contained in the genetic code. The GC content of the underlying nucleotide sequence is itself not unusual. This means that Goddard is unlikely to have emerged from a “random” stretch of ncDNA since it would not have reflected the bias in the amino acid frequency distribution of its composition.

**Figure 3:**
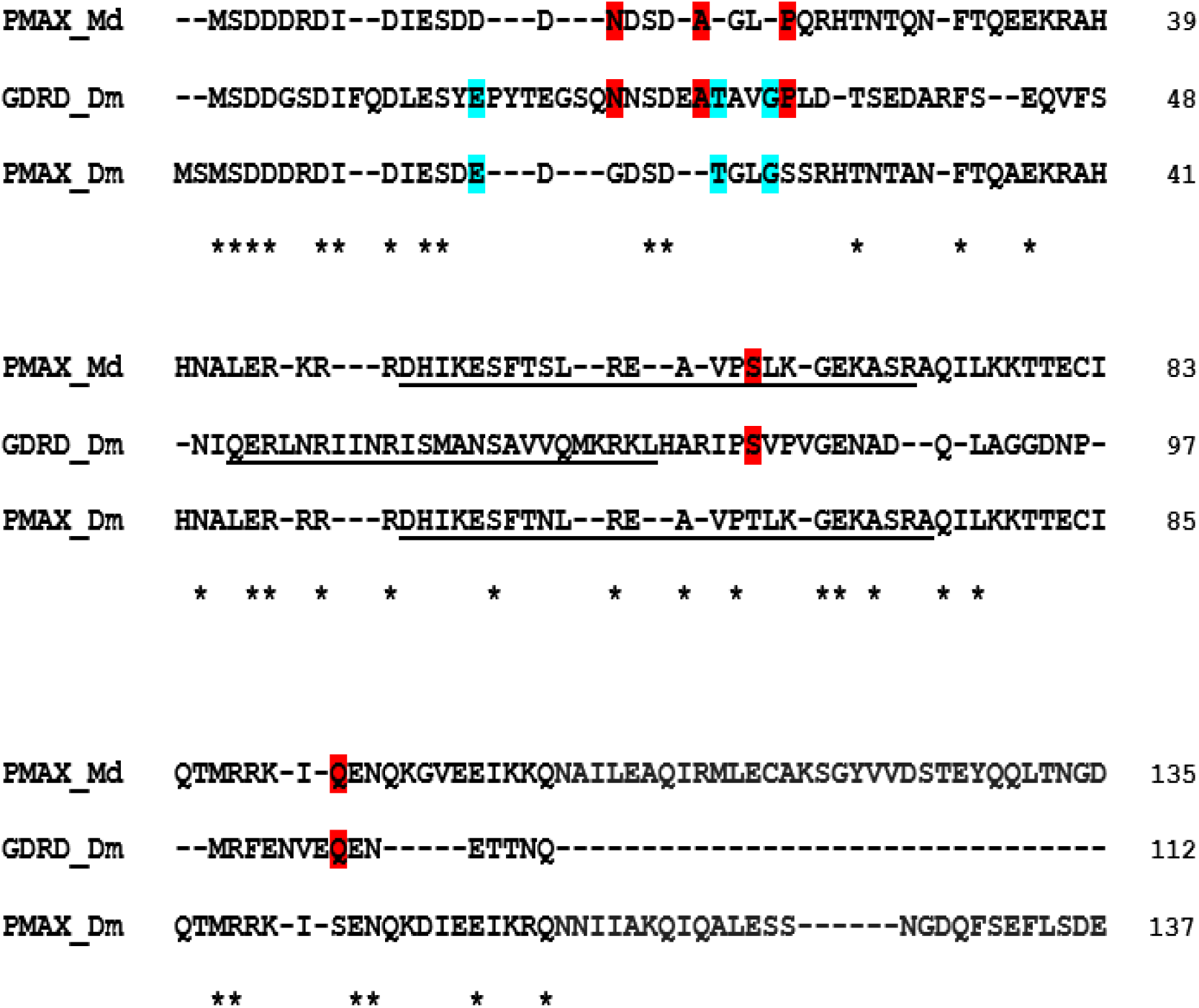
The alignment of Goddard in Drosophila melanogaster (Dm) alongside Protein MAX in the same species and with Musca domestica (Md). Not all the Protein MAX sequences are displayed. Asterisks denote shared residues in all three sequences. Underlined are those residues that are found to be the most structurally aligned between Goddard and MAX. Shaded areas denote residues shared between Goddard and just one of the MAX genes. As can be seen, some residues present in Goddard (A29, T30, G33) are not present in one of the MAX genes but are found in the other as would be fully expected amongst diverging paralogs.

Interestingly, Protein MAX in *D. melanogaster* itself is only about 28% identical, and there seems to have been an internal amplification of the first two residues (MS) in it and other Drosophila species. In *C. capitata,* MAX is just 24% identical with Goddard. However, the sequences serve to fill in for residues present in Goddard but missing in the others and gaps may not just due to deletions in both genes. The C-terminus of Protein MAX, containing the localization signal, that facilitates the transporting of the protein from the cytoplasm into the nucleus, is also absent in the alignment implying that there has been a premature truncation. This means that it is unlikely Goddard would perform the same DNA-binding functions that MAX does, and that whatever function it is involved with must be different; MAX itself is observed to function both as a repressor and activator where it is located mainly in the nucleus (Kato et al. 1992).

The 5’ untranslated regions, moreover, as shown in Suppl Fig 4, reveal the presence of a large bulk deletion in the respective sequence for Goddard when they are aligned. Aside from this, there is a high degree of identity in places, especially upstream of this deletion, but this alignment is still interrupted with many gaps and BLASTN does not conclude it is significant. It is unclear if co-regulation mediated by promoter and 5’ UTR sharing (Yang et al 2008) occurs between Goddard and its host in Omega; transcriptional interference could explain some of the differences in expression (Lee and Chang 2013). The existence of an independent basal promoter is inferable but certainly not confirmed by a transgenic experiment. One possibility is that enhancers located in the UTRs could confer the ability to act as or interact with other promoters, not just Omega’s, and so effect transcription (Kowalczyk et al 2012; Lee and Wu 2006).

In terms of structure, both Goddard and MAX are computationally predicted to contain alpha helices with stable folds. According to Lange et al. (2021), a second, shorter helix is also predicted close to the N-terminus of the former, but with lower confidence. Figures 4a/4b display default conformations for both proteins. Goddard, thus, may either assume a helix-loop-helix, as with MAX, or a loop-helix-loop structure instead. When structurally aligned, there is great similarity with an RMSD of 2.37 Å, reflective of at least 55 residues that can be directly juxtaposed. The structural alignment is shown in Suppl Fig 5. In terms of protein aggregation, both Goddard and MAX are predicted to be almost exactly disordered at 60%. Aggregation can lead to reduced DNA binding and impaired transcriptional activity. Moreover, any intrinsic disorder allows for greater versatility in protein interactions (Dyson and Wright 2005).

**Figure 4:**
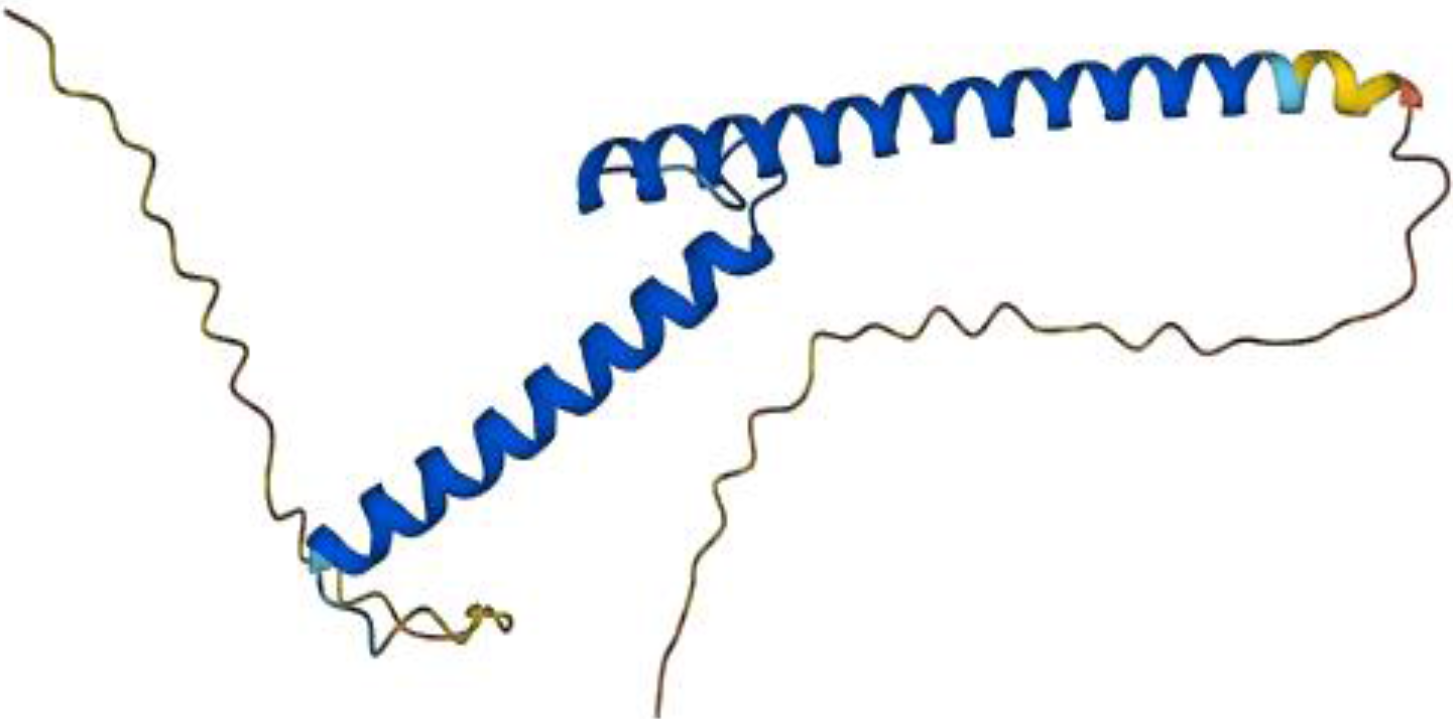
The predicted helix-loop-helix structure of Protein MAX in Drosophila.

**Figure 4b:**
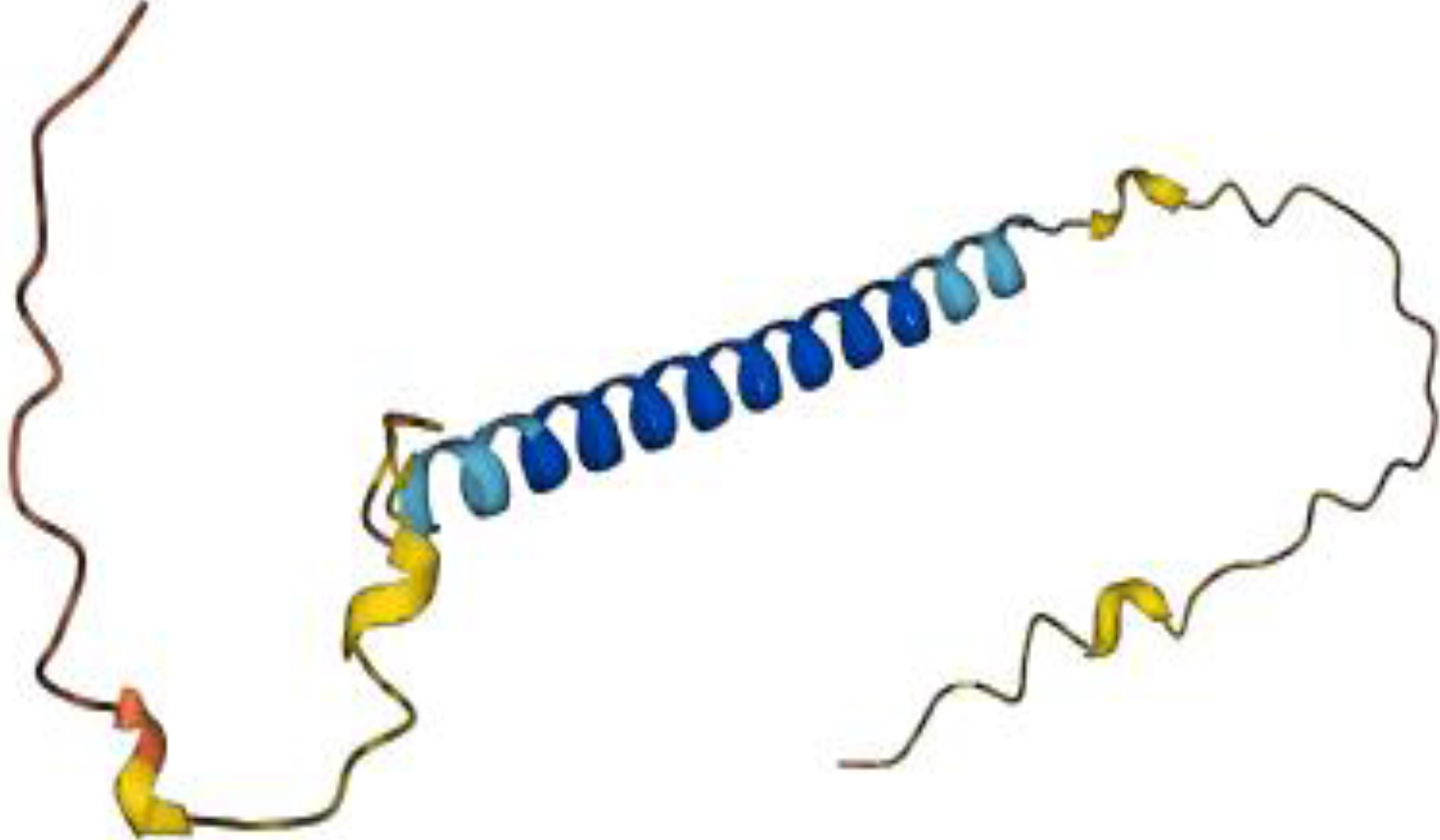
The predicted loop-helix-loop structure of Goddard in Drosophila. A second helix may exist but confidence for this is not high. It could, therefore, resemble MAX even more so.

Further computational tests, using both composition and PSSM approaches, show that Goddard and MAX in Drosophila may *both* be RNA-binding. Indeed, RNA binding by DNA-binding proteins is found to be unusually commonplace (Cassiday and Maher 2002): various transcription factors are now proposed to form functional interactions with mRNA to facilitate the precise regulation of gene expression (Holmes et al 2020). The basis for promiscuous protein-nucleic acid interaction is not addressed here. Goddard is also predicted to form heterodimers, like MAX, a possibility admitted by (Lange et al 2021), even if observations supporting this are lacking. Spermatogenesis in Drosophila is highly reliant on mechanisms of post-transcriptional regulation of gene expression, as facilitated by RNA-binding proteins (Sutherland et al 2015); the knocking out of TULP2, an RNA-binding protein, leads to male sterility in mice (Zheng et al 2021). Suppl Tables 5 and 6 display the respective RNA-binding residues for both genes whilst Suppl Tables 7-8 reveal the predicted list of functions that include RNA binding for MAX. Goddard may be like some SOX members that interact with RNA to regulate genes expressed within the testes (Kiselak et al. 2010; Zhang and Hou 2021).

Protein MAX has a low tissue-specificity pattern of expression, but Goddard is very much testis-specific. It is widely acknowledged from numerous studies that, within *Drosophila* at least, retrogenes are more prone to have a testis-specific expression whereas older duplicates are much more often ubiquitously expressed or specifically expressed in other somatic tissues (Zhang and Zhou 2019; Assis and Bachtrog 2013; Bai et al. 2008; Guschanski et al. 2017). According to Kumar et al (2009), whilst reporting the findings of Assis et al (2008), “the vast majority of nested intronic genes emerge through the insertion of a DNA sequence into an intron of a preexisting gene”. Retrogenes have, moreover, been found both within introns and intergenic regions alike (Vinkenbosch et al. 2006). Given Goddard’s location within an intron of another gene, it is likely that the gene is itself a retroposed and truncated copy of MAX, which itself contains 4 introns that are not present in the former; the origin of any internal gene is classified as being retrotransposition when it is itself intronless. The de novo hypothesis offers no explanation to show how just part of the sequence of an intron became its own transcript to independently recruit the polymerase and be translated, let alone becoming a protein required for male fertility in most fruit flies. Goddard may, instead, be an example of some degree of functional partitioning among paralogs.

### BSC4: de novo birth or gene fission?

BSC4 is a 131 aa protein encoded within *S cerevisiae* that is estimated to be about 100 million years old (Casci 2008). Its name indicates that it bypasses stop codons present in the coding sequence. Orthologous sequences in other yeast species are non-coding, and it was presented as one of the first examples of how an entirely new gene, not just part of one, could potentially arise from an already transcribed non-coding region. The gene is believed to have obtained an ORF of viable length through a series of point mutations. BSC4 expression is up-regulated during the stationary phase of the cell cycle, and it may be involved in the DNA repair pathway. It might, therefore, contribute to the robustness of cells in a nutrient-poor environment (Cai et al. 2008). Curiously, BSC4 is itself one of several stop codon read-through genes that avoid premature termination despite interruptions in the reading frame (Namy et al. 2003).

According to Weisman (2022), a ribosome profiling dataset performed by McManus et al. (2014) of the closest relative, *S. paradoxus* reveals very low but non-zero protein expression, suggestive of ancestral “leaky” translation. Indeed, one possible explanation is that the peptide status of BSC4 was simply lost or reduced in other yeast lineages except for *S. cerevisiae*. Another is that, like FLJ33706, it is a pseudogene that has been revived in just one yeast lineage alone. The gene has no homolog in any fungal species, supporting its de novo status, although Cai et al. (2008) admit that horizontal gene transfer, or very high divergence in an ancestral gene, is a possibility. As in the previous examples, the N-terminus of BSC4 consisting of the first 22 residues, was BLASTed against the proteome of *S. paradoxus* to see if any potential hits of interest were produced. Again, none were better than the threshold for significance, but one of the hits displayed a marked cluster of shared residues that warranted further examination. The presence of gaps and amino acid repeats can account for a relatively high E-value. Unlike in the previous examples, the N-termini are out of position. When the entire BSC4 sequence is aligned with the 439 aa protein, Vacuolar sorting protein 38 (Vps38), it is clear that the N-terminus of the former corresponds to most of the latter’s C-terminus. By itself, this finding contradicts the assumption made about align-ability, except that there are examples of gene fission (Wang et al. 2004; Paseck et al. 2006) whereby the duplicate of a gene either splits in two to form separate genes, or where one part is retained, and the other is lost. Since many genes are modular, being comprised of discrete units, a fission could still prove to be viable.

Both BSC4 and Vps38 contain just the one exon and, overall, there is a 27% sequence identity with the corresponding sequence for the latter in *S. cerevisiae*. This would thus place it within the twilight zone of homology. The alignment is shown in Figure 5. N8 in BSC4, however, is absent in Vps38 for cerevisiae but present in paradoxus. Not all the C-terminus is present in BSC4 which, again, indicates a premature truncation is likely. If Vps38 is the precursor sequence, it appears that a valine codon was mutated to become a novel start site. Interestingly, the initial methionine codon in BSC4 is also valine (GTG/GTA) in the orthologous non-coding sequences of the other yeast species. In terms of amino acid composition, BSC4 is much more alkaline than the C-terminus of Vps38. It seems to have preserved key lysine residues, along with much less frequent tyrosine and phenylalanine amino acids that may influence catalytic activity. A chi square test for the amino acid composition of BSC4 produced a P-value < 0.01. Lysine and isoleucine are over-represented but the frequency of glycine is low. The GC content is only slightly lower than average, and thus does not explain this distribution.

**Figure 5:**
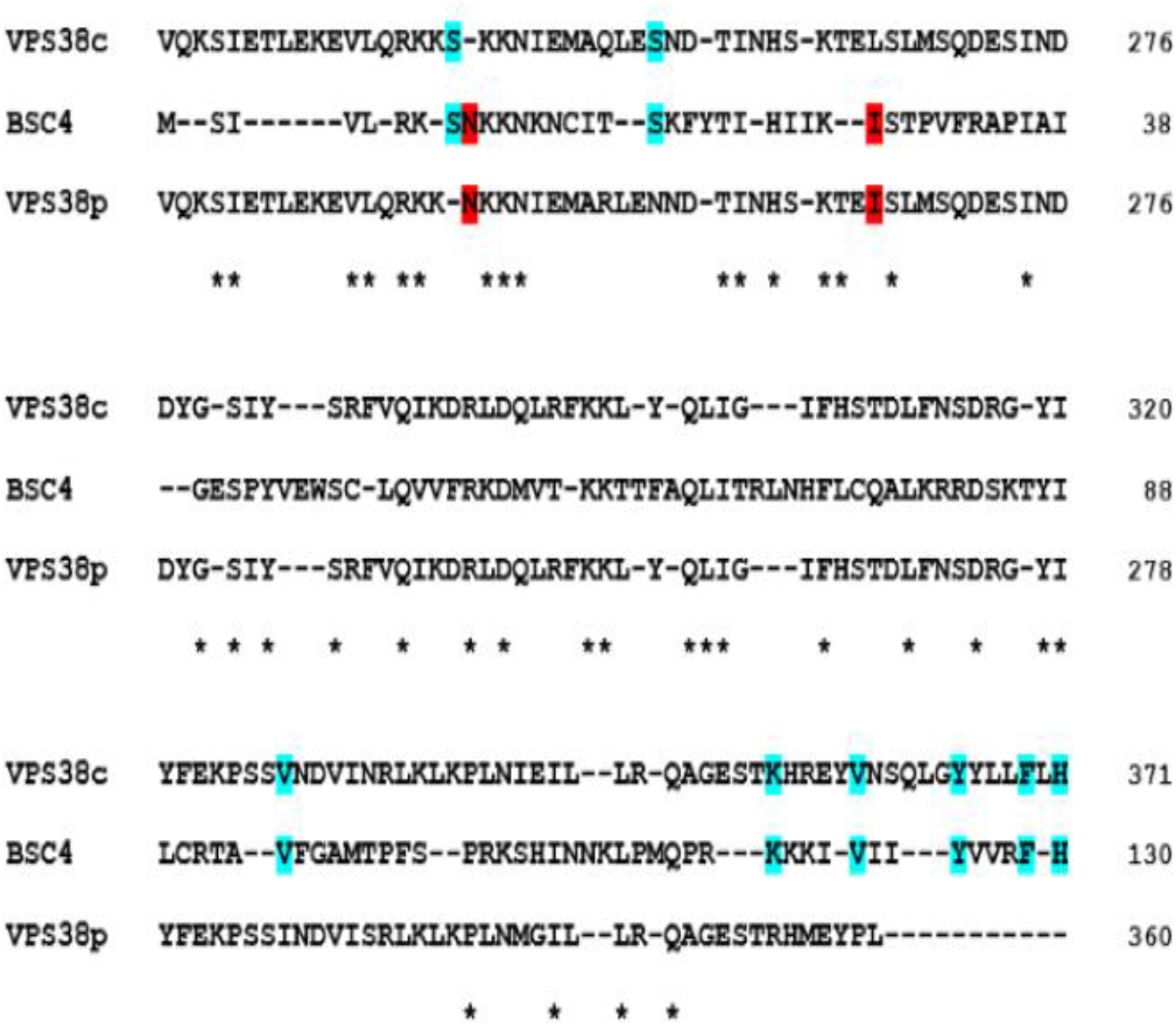
The alignment of BSC4 and the corresponding sequence at the C-terminus of VPS38c (S. cerevisiae) and VPS38p (S. paradoxus). The latter two start with valine exactly as predicted in non-translated orthologs of BSC4 within other yeast species. Asterisks show shared residues that are identical in all three protein sequences. Shaded areas denote those residues found in BSC4 but only in one of the yeast orthologs as well. The S7 and N8 sites in BSC4 are present in one but not the other of the Vps38 sequences, which is common in evolutionary divergence.

The structure of the BSC4 protein is predicted by Bungard et al. 2017 to have a single rudimentary fold consisting mostly of beta sheet conformation. It is also stable with few residues predicted to be disordered. By contrast, Vps38 is predicted to consist mostly of alpha helices, as can be seen in the secondary sequence shown in Suppl Fig 6, with a higher level of intrinsic disorder. About five residues, however, also appear to form beta sheet. This would imply there is no significant structural homology between them although some models indicate the presence of two alpha helices in BSC4, as in Suppl Fig 7, and an RMSD of 3.56 Å was calculated. Even though structure tends to diverge less than sequence does (Williams and Lovell 2009), conformational transitions from alpha helix to beta sheet are actually quite common (Ding et al. 2017).

Of particular interest is the function of Vps38 and how it relates to BSC4’s possible role in the signal pathway for DNA damage. The gene binds with both Vps30 and Vps34 to promote production of phosphatidylinositol 3-phosphate required to stimulate kinase activity (Liu et al 2018). It is involved in facilitating both vacuole transport and autophagy to preserve cellular homeostasis. Phosphatidylinositol-3 kinases are known to play an important role in relaying DNA damage signaling from break sites (Khan et al. 2019). Indeed, autophagy may itself have evolved as a quality control system that responds to a wide range of stress conditions that include DNA damage (Dotiwala et al. 2012; Eliopoulos et al. 2016; Ajazi et al. 2021). BSC4 would appear to interact with DUN1 (Pan et al. 2006), a kinase that is required for the damage-induced transcription of DNA repair genes (Zhou and Elledge 1993; Yam et al. 2020), and both Vps38 and DUN1 are included in the same interaction network by Kramer et al (2017). Functional prediction (Suppl Tables 9-10) indicates that that both BSC4 and Vps38 are involved in catalytic activity and cytoskeletal binding, the latter known to have a role in autophagy (Kast and Dominguez 2017). Subcellular localization prediction shows that BSC4 is related to genes localized to the mitochondrion or Golgi body, including DUN1 and Vps38, and a mitochondrial transfer peptide is detected at the N-terminus and is displayed in Suppl Fig. 8. There is, thus, sufficient sequence identity and functional homology for Vps38 to be considered a potential homolog for BSC4. In such a scenario, the copy of the C-terminus of a gene was preserved by selection in one yeast lineage.

### AFGPs from non-genic DNA in cod: a non-sense story

By far the most striking claim made about de novo origination is the purported creation of antifreeze glycoproteins in cod species out of “junk” ncDNA. Zhuang et al. (2019) offer an account that describes how a functional and essential gene was formed stepwise entirely in the absence of natural selection, only later improving once the sequence had been transcribed and translated. It appears to validate the proto-ORF model (Guerzoni and Mclysaght 2015) whereby an ORF existed before the regulatory element to activate transcription was gained, in this case via a putative translocation. Previously, Baalsrud et al (2018) had suggested that afgps in codfish likely arose from ncDNA because, among other factors, their GC content is unusually high and because of their intrinsic structural disorder – consistent with observations made about de novo genes. However, they also acknowledge that, “the alternative scenario is that afgp evolved from a pre-existing protein encoding gene so diverged from afgp to not be recognized by BLAST.” Even so, they consider such rapid divergence to be unlikely. The scenario envisaged is highly dependent on the “fortuitous” presence of a Kozak sequence, a signal peptide and a glutamine-rich propeptide contained within “latent coding exons”. It also requires that the 3’ end of the gene was derived from a non coding sequence excessively prone to intragenic recombination (Zhuang and Cheng 2021). Previous research into the origins of antifreeze proteins had focused on duplicate genes as the likely precursors to antifreeze proteins (Deng et al. 2010; Chen et al. 1997). The reasons given for offering a de novo interpretation are as follows: i) there are no meaningful homologs to any part of the gadid afgps to hint at ancestry and, in particular, neighboring loci bear no resemblance to them ii) orthologous non coding sequences in some basal species resemble the coding sequences of the afgp genes. The researchers make an *a priori* assumption, namely that the encoded “Thr Ala-Ala” amino acid triplets, that are essential to ice-binding functionality, must have arisen through the successive and exclusive internal duplication of a specific element. The model of de novo birth proposed also entails various single nucleotide substitutions and a frameshift to produce a gene replete with both introns and UTRs. The process outlined seems to be congruent with the hypothesis of constructive neutral evolution (CNE), first proposed by Stolzfus (1999), whereby neutral changes can account for the development of complexity in biology (Muñoz-Gómez et al 2021).

There also appears to be some confusion as the authors hypothesize that, “upon the onset of selective pressure from cold polar marine conditions, duplications of a 9-nt ancestral element in the midst of the four GCA-rich duplicates occurred.” However, they appear to be explaining a stage relating to what happened *before* the gain of the promoter allowing the sequence becoming transcribed, not just *after* it had occurred. This is most probably an oversight rather than a failure to understand that the gene would only have been useful once it had been translated. As has been seen, moreover, homologous genes do not have to be positioned closely to each other. If the authors can imagine that the sequence was transposed to a region containing a promoter, it is unclear why they see the absence of sequence similarity in neighboring genes as significant. The reconstruction is problematic because the stepwise changes to the sequence appear to have all survived random drift, whilst not suffering any potentially degrading mutations, despite not having been selected. The authors even claim this did, in fact, happen to afgp genes in some species, resulting in their pseudogenization. Moreover, in this initially selection-free ORF-first scenario, we might justifiably ask why similar proto-ORFs are not found in organisms that do not living in the icy colds?

As in the previous examples, the N-terminus consisting of the first 21 residues of one of the smallest antifreeze glycoproteins identified by Zhuang et al. (2019), namely AFGP2 in the Atlantic cod, was BLASTed against all gadids to see if there were any notable hits. This gene may represent something resembling the original afgp of the species. Aside from the other afgps, several interesting hits with an E-value < 0.1 were returned, all aligned with AFGP2 at their respective N-termini. One of them was for a 275 aa apolipoprotein A-II. This warranted attention given that its product transports hydrophobic lipids in the blood plasma, as antifreeze proteins do, except that the latter bind to ice crystals. Lipoproteins, in fact, have a functional and evolutionary relationship with antifreeze proteins (Gauthier et al 2008), including the 125 aa GmAFPIV in the Pacific cod that serves a dual function as a antifreeze and lipoprotein (Mao et al. 2018). Indeed, Cheng (1998) cites them as evolutionary precursors to Type IV afps in other piscine species, even though the sequence identity is just 22% (Deng et al. 1997), since their polar surfaces can bind with ice. Even Antarctic bacteria have acquired an antifreeze capability through utilizing a lipoprotein (Yamashita et al 2002).

The coding sequence of ApoA-II, in cod and zebrafish alike, is interrupted by two introns as is the case in all the afgps. Figure 6 shows the aligned sequence of AFGP2 in the Atlantic cod, as well as the Arctic cod, along with the apolipoprotein in the former species and an ortholog in an outgroup species, D. rerio. Suppl Fig. 9 also shows the aligned 5’ UTRs of AFGP2 in the Atlantic cod and its possible precursor. There is a notable identity between the afgps and lipoproteins, with an amino acid identity of up to 32%, mostly because of the highly similar signal peptides. As stated above, Chen et al. (1997) identified a trypsinogen gene as the precursor of afgps in notothenioids on account of its signal peptide. The zebrafish apolipoprotein sequence fills in for certain residues, with 8 sites uniquely matching those in one or both afgps. In total, as many as 52 of the aligned 124 residues in AFGP2 in the Atlantic cod have identical sites in the apolipoproteins displayed. As in the previous examples, a premature truncation has likely happened. The gene coding for the protein in the cod species, including UTRs, is also rich in (G)CA, the presence of which is seen as the basis for alanine in the repeating elements. In other afgps, these repeats are more numerous, implying that internal amplification of this motif was a factor in the sequence expansion. However, it may not have been required for the initial state of the ancestral afgp. Other than the notable difference in alanine frequencies, the amino acid composition is very similar.

**Figure 6:**
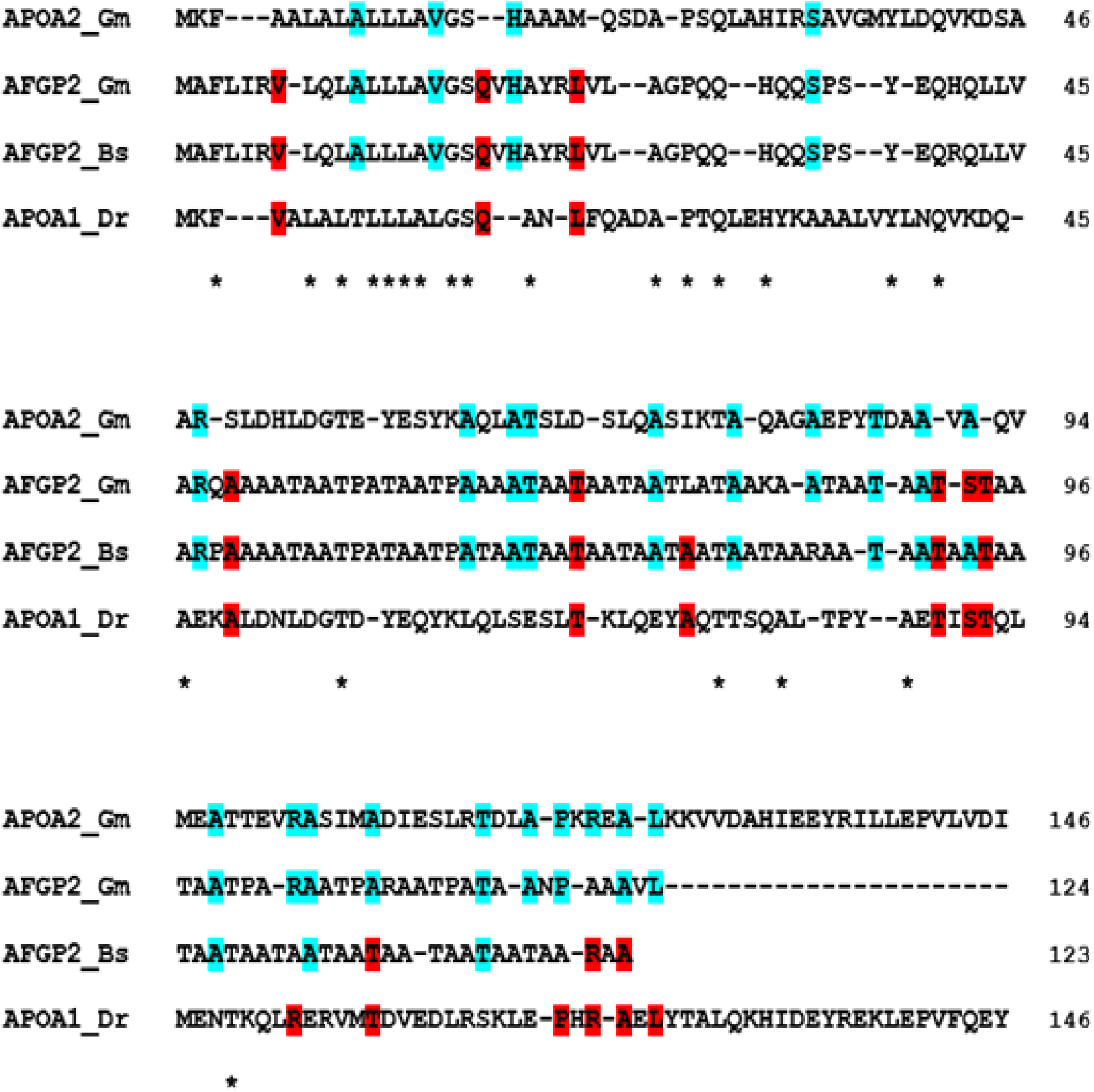
The alignment of AFGP2 in the Atlantic cod (Gm) and Polar cod (Bs) along with apolipoprotein (A-II) in the Atlantic cod, top sequence, and apolipoprotein (A-I) in the zebrafish (Dr) at the bottom. Not all of the longer apolipoprotein sequences are displayed. Also, the entire sequence of the Polar cod afpg is not shown since a number of repeating ‘TAA’ trimers are not present in the Atlantic cod but the protein terminates with “VL” as with its ortholog. Asterisks denote residues shared among all proteins. Shaded areas denote residues shared between lipoproteins and one but not both afgps. Although the lipoprotein in Atlantic cod is closer to the afgp sequence of the same species, a couple of key sites in the afgp, namely V6 and Q18, are found exclusively in that of the zebrafish and these may prove to be primitive.

In terms of its secondary structure, the apolipoprotein appears to be much more consistent in form than the antifreeze protein, and is made up largely of alpha helix, as seen in Suppl Fig 10. However, Suppl Fig 11, indicates the presence of stable helices in the latter’s tertiary structure. A prediction of lipid-binding propensity for both proteins, as displayed in Suppl Fig 12, shows that the antifreeze protein has retained vestiges of ancestral functionality, at least at the N-terminus. Research has shown that signal peptides can interact directly with the hydrocarbon region of lipids (Harrington et al 2014). This comports with the idea that af(g)ps bind to lipids because of the similarity of helix bundle to apolipoproteins and so could be pre-adapted to having ice as the ligand (Ghalamara et al 2022). A tertiary structural alignment produces an RMSD of 3.02 Å. Even if the afgps did emerge from a non-coding sequence, as claimed, it was likely an apolipoprotein pseudogene rather than an arbitrary piece of DNA that happened to contain the constituent components of an antifreeze protein. This would account for the “substantial variation” between species and a “lack of functional constraint”. The inferred frameshift would have restored the original reading frame of a pre-existing gene sequence, whilst the substitutions (mainly for alanine) paved the way for the evolution of ice-binding function that perhaps was present already. This is a far more elegant account, and entirely congruent with evolutionary theory, than a speculative “something out of nothing” story that is highly dependent on serendipity.

## Analysis

The previous section discovered that, contrary to previous claims, meaningful and plausible homologs exist for the four de novo candidate genes considered, indicating that duplication and divergence could potentially be a valid alternative explanation. A comparative account of these is necessary to summarize and evaluate their relative strengths and weaknesses by using several features/metrics relating to gene ontology.

In Tables 1-2, displayed below, there are five key metrics namely i) sequence, ii) structure, iii) functionality, iv) composition and v) deployment listed with respective sub-metrics totaling twelve. These have all been described in detail, except for the correlation coefficient for amino acid frequencies (composition) that is found in the Suppl Dataset for each gene. This is relevant for assessing compositional similarities among potential homologs whereas the calculated P-value, with respect to expected frequencies in the genetic code, relates to the likelihood that the distribution of amino acids is reflective of a possibly random stretch of underlying DNA. This allows for a transparent assessment of the likelihood that the gene did arise de novo or through a duplication event instead. In the case of assessing the specific merits of de novo gene birth, three categories (unlikely, possible and likely) were used when evaluating the relative plausibility. For example, a very low E-value would be regarded as unlikely in a scenario where a gene is claimed (perhaps incorrectly) to have no homologs. But the outcome is not a zero-sum in as much as both claims can be equally strong or weak.

**Table 1:** The summary for the alternative scenarios in FLJ33706 and Goddard with respect to duplication or de novo origination. This is based on the results obtained.

| Gene Feature | De novo candidate | Potential homolog | De Novo Status |
| --- | --- | --- | --- |
| Name/Origin | FLJ33706/c20orf203 | ZNF678.1 (5' end) | lncRNA |
| **Sequence** |  |  |  |
| *E-values* |  | Low (whole and part) | Unlikely |
| *AA identity* |  | Relatively Low to Fair | Possible |
| *Organization* | Multi-exon | Multi-exon | Possible |
| **Structure** |  |  |  |
| *Conformation* | Alpha helix (Krab domain) | Alpha helix (Krab domain) | Possible |
| *RMSD* |  | Relatively Low | Possible |
| *Disorder level* | Partly disordered | Not disordered | Likely |
| **Functionality** |  |  |  |
| *Class* | Transcription factor | Transcription factor | Unlikely |
| *Binding affinity* | DNA-binding | DNA-binding | Unlikely |
| **Composition** |  |  |  |
| *Correlation coefficient* |  | Weak relationship | NA |
| *P-value (genetic code)* |  | Very significant | Unlikely |
| **Deployment** |  |  |  |
| *Expression* | Brain and gonads | Brain,thyroid and gonads | Possible |
| *Localization* | Nucleus | Nucleus | Unlikely |
|  | **OVERALL** | Strong similarity | Unlikely |
| Name/Origin | Goddard | Protein MAX | Intron |
| **Sequence** |  |  |  |
| *E-value* |  | Low (part) | Possible |
| *AA identity* |  | Relatively Fair | Possible |
| *Organization* | Single exon | Multi-exon | Likely |
| **Structure** |  |  |  |
| *Conformation* | Helix-Loop | Helix-Loop-Helix | Possible |
| *RMSD* |  | Low (full length) | Unlikely |
| *Disorder level* | Largely disordered | Largely disordered | Likely |
| **Functionality** |  |  |  |
| *Class* | Regulatory protein | Regulatory protein | Unlikely |
| *Binding affinity* | DNA/RNA binding | DNA/RNA binding | Unlikely |
| **Composition** |  |  |  |
| *Correlation coefficient* |  | Fair relationship |  |
| *P-value (genetic code)* |  | Very Significant | Unlikely |
| **Deployment** |  |  |  |
| *Expression* | Testes | General incl the testes | Possible |
| *Localization* | Axoneme; Cytosol | Nucleus; Cytoplasm | Unlikely |
|  | **OVERALL** | Strong similarity | Unlikely |

**Table 2:** The summary for the alternative scenarios in BSC4 and AFGP2 with respect to duplication or de novo origination. As above, this is based on the results obtained.

| Gene Feature | De novo candidate | Potential homolog | De Novo Status |
| --- | --- | --- | --- |
| Name/Origin | BSC4 | VPS38 (3' end) | lncRNA |
| **Sequence** |  |  |  |
| *E-value* |  | Fair (part) | Likely |
| *AA identity* |  | Relatively Fair | Possible |
| *Organization* | Single exon | Single exon | Likely |
| **Structure** |  |  |  |
| *Conformation* | Beta sheet (mainly) | Alpha helix (mainly) | Likely |
| *RMSD* |  | Relatively Fair | Possible |
| *Disorder level* | Not disordered | Partly disordered | Possible |
| **Functionality** |  |  |  |
| *Class* | Autophagy gene | Autophagy gene | Unlikely |
| *Binding affinity* | Protein binding | Protein binding | Likely |
| **Composition** |  |  |  |
| *Correlation coefficient* |  | Weak relationship | NA |
| *P-value (genetic code)* |  | Significant | Unlikely |
| **Deployment** |  |  |  |
| *Expression* | NA | NA | NA |
| *Localization* | Mitochondrion | Mitochondrion/Vacuole | Unlikely |
|  | **OVERALL** | Fair similarity | Possible |
| Name/Origin | AFGP2 | APOA2 (5' end) |  |
| **Sequence** |  |  |  |
| *E-value* |  | Low (part) | Possible |
| *AA identity* |  | Medium | Unlikely |
| *Organization* | Multi-exon | Multi-exon | Possible |
| **Structure** |  |  |  |
| *Conformation* | Loop with helix | Mainly alpha helix | Possible |
| *RMSD* |  | Relatively Low | Possible |
| *Disorder level* | Not disordered | Largely disordered | Likely |
| **Functionality** |  |  |  |
| *Class* | Anti-freeze glycoprotein | Transport protein | Possible |
| *Binding affinity* | Ice crystals; (lipids) | Lipids; (water) | Possible |
| **Composition** |  |  |  |
| *Correlation coefficient* |  | Strong relationship | NA |
| *P-value (genetic code)* |  | Significant (caveat) | Possible |
| **Deployment** |  |  |  |
| *Expression* | Pancreas (mainly) | Liver (mainly) | Possible |
| *Localization* | Extracellular | Extracellular | Unlikely |
|  | **OVERALL** | Fair similarity | Possible |

Table 3, moreover, examines the likelihood that the mechanisms/events responsible for the evolutionary origination and propagation of the genes is consistent with those for de novo birth or duplication and subsequent divergence. Other metrics include creation of the underlying substrate, transcription gain, and that of the introns and UTR, to support the gene’s stability and regulation. One aspect not mentioned in the analysis concerns the feasibility of fixation broken down into adaptive utilization and any pleiotropic effects. The former refers to the ability of a new gene to be utilized in such a way that contributes to improving fitness and so being favored by natural selection. This would typically involve how the gene is incorporated into biochemical pathways and deployed so that it readily confers a survival advantage. In the case of the antifreeze glycoprotein, this would be readily apparent since secretion into blood plasma would have an immediate effect. For the others, a route to fixation is less clear. Heames et al (2023) report that de novo candidate genes are found to be much more soluble than random library sequences that nonetheless mirrored their composition. Pleiotropic effects, on the other hand, refer to the extent to which a gene potentially has multiple phenotypes: pleiotropic conflict is expected to reduce the efficacy of selection by limiting the fixation of beneficial mutations through adaptation (Fraïsse et al 2019). If a gene’s expression is not tightly regulated, as might be the case for a new de novo gene, it could be antagonistically pleiotropic thereby limiting its adaptive utility. Both sub-metrics assume that fixation is achieved largely through selective pressures whereas drift is either ancillary or has no effect at all. The strength of the latter, of course, depends on population size not just the environment (Kimura 1964).

**Table 3:** The summary for the inferred mechanisms of gene production in all four genes with respect to the alternative scenarios of duplication and de novo origination.

| Gene Feature | De Novo Candidate | Potential Homolog | De Novo Status |
| --- | --- | --- | --- |
| Name/Origin | FLJ33706 | ZNF678 (5'end) | lncRNA |
| **Substrate creation** | Intergenic region | Retro-duplication | Likely |
| **Transcription gain** | Open chromatin | Open chromatin | Possible |
| **ORF generation** |  |  |  |
| *Length* | Stop site removal | Stop site removal | Likely |
| *Structural motifs* | Fortuitous | Innate to paralog | Unlikely |
| **Intron/UTR gain** | Alu insertions | Alu re-insertions | Possible |
| **Fixation** |  |  |  |
| *Adaptive utilization* | Unclear | Probably redundant | Possible |
| *Pleiotropic effect* | Different traits | Not a major issue | Possible |
|  | **OVERALL** |  | Possible |
| Name/Origin | Goddard | Protein MAX |  |
| **Substrate creation** | Intronic region | Retro-duplication | Likely |
| **Transcription gain** | Unclear | Enhancers present | Possible |
| **ORF generation** |  |  |  |
| *Length* | Avoidance of stops | Truncation | Possible |
| *Structural motifs* | Fortuitous | Already present | Possible |
| **Intron/UTR gain** | Uncertain UTR | Already present | Possible |
| **Fixation** |  |  |  |
| *Adaptive utilization* | Uncertain | Uncertain | Possible |
| *Pleiotropic effect* | Very limited | Tissue-specific | Unlikely |
|  | **OVERALL** |  | Possible |
| Name/Origin | BSC4 | VPS38 (3'end) | lncRNA |
| **Substrate creation** | Intergenic region | Duplication/Fission | Likely |
| **Transcription gain** | Transcribed region | Transcribed region | Likely |
| **ORF generation** |  |  |  |
| *Length* | Stop codon bypass | Stop codon bypass | Possible |
| *Structural motifs* | Unclear presence | Basic form present | Possible |
| **Intron/UTR gain** | Uncertain UTR | Already present | Possible |
| **Fixation** |  |  |  |
| *Adaptive utilization* | Unclear integration | Part of network | Unlikely |
| *Pleiotropic effect* | Unknown | Mainly restricted | Unlikely |
|  | **OVERALL** |  | Possible |
| Name/Origin | AFGP2 | APOA2 (5' end) |  |
| **Substrate creation** | Amplification | Duplication | Unlikely |
| **Transcription gain** | Translocation | Promoter present | Unlikely |
| **ORF generation** |  |  |  |
| *Length* | Incremental accretion | Truncation | Unlikely |
| *Structural motifs* | Fortuitous presence | Basic form present | Possible |
| **Intron/UTR gain** | Chance frameshift | Already present | Unlikely |
| **Fixation** |  |  |  |
| *Adaptive utilization* | Immediate gain | Immediate gain | Likely |
| *Pleiotropic effect* | Some potential | Some potential | Possible |
|  | **OVERALL** |  | Unlikely |

The summary tables (1-3) indicate that Goddard is the most likely to have a homolog and been created by gene duplication with FLJ33706 and AFGP2 not far off. In the case of the latter, the inferred generation of the ORF from an arbitrary stretch of recombination-prone DNA, and subsequent incremental accretion, is adjudged as being implausible, whereas the ORF would have been inherited intact from an apolipoprotein paralog that was truncated. The P-value for the composition includes a caveat because of the highly repetitive nature of the sequence and GC-bias. BSC4, however, is found to be the most marginal with the overall likelihood little different than for a de novo origin. The evaluation was conducted to be as generous to the de novo hypothesis as possible. Whilst for some sub-metrics, a de novo origin can be argued, the overall breakdown falls in favor of a traditional model of gene birth as the best explanation. The mechanisms responsible, including its fixation in the population, appear to be more appropriately attributed to duplication than to a de novo creation. Both FLJ33706 and BSC4 are suggested as having evolved from lncRNAs, which may have served an intermediate function, whereas Goddard and AFGP2 would had to have acquired their own transcriptional regulation. There is, nevertheless, sufficient evidence detracting from the proposed idea that the candidates were recruited from stretches of ncDNA, that were never coding, and that they do have plausible homologs in genes whose copies they were created from. The approach, as used here, could be applied for other de novo candidates should potential homologs be identified for them.

Unfortunately, the summary tables do not include a precise calculated probability of either scenario across the various metrics examined which might have been useful. Such quantification, although provided within the discrete results displayed in the supplementary tables and dataset, are not translatable for the comparative analysis; the evidence presented is cumulative but not additive or multiplicative. A hypothetical “coefficient of similarity” cannot be determined, and, in any case, homology is not a continuous variable but is a boolean value. Statistical analysis alone is insufficient, moreover, especially where the signal of homology is weak using standard detection methods, and so needs to be supported by a valid understanding of biological reality.

## Discussion

Although once not seriously considered as being likely (Kaessmann 2010), the de novo origination of entire genes from non-genic DNA has since become a dogma in the study of molecular evolution (Cherezov et al. 2021). This would apparently defy the expectation of Jacob (1977) namely that, “the probability that a functional protein would appear de novo by the random association of amino acids is practically zero.” This has resulted in extensive searches for candidate de novo genes, and evolving “proto-genes”, in all species (Parikh et al. 2022). Part of the problem here has been that researchers in the field have relied too heavily on similarity search tools, like BLAST, when trying to find out or rule out potential antecedents. But they are not perfect, as has been demonstrated, and many homologs may be missed because the algorithm struggles to pick up on related sequences held in a database that have been subject to significant change. Processed pseudogenes tend to evolve more rapidly than their functional paralogs (Tutar 2012), and if reactivated their relationship to other genes could be quite hard to determine. Even without gaps, sequences that have diverged substantially will likely either be missed or returned with E-values much worse than the default threshold. A tendency to see only statistically significant matches as relevant can preclude further investigation where there is, at least, a suspicion that two sequences could be related owing to one or more shared features, however minor, and so false negatives occur. With the benefit of hindsight, and also greater domain knowledge, the four homologs presented perhaps all should have been suspected prior to a BLAST search which served to identify possible related genes.

One definition for d*e* novo genes is that they are protein-coding genes having no clear homology to previously existing protein-coding genes (Reinhart et al. 2013). The other, of course, is that they arise from non-coding DNA. The first principle would appear odd since it implies that the definition of a de novo gene is one for which a standard methodology of homology detection is insufficient. The failure to detect any homologs has become one of the reasons offered to proclaim the existence of a candidate de novo gene. But the absence of such a homolog might be due to the incompleteness of genomic databases, although this is now becoming far less of an issue. As seen here, there are other reasons as to why homology detection can and will consistently fail. The totality of the evidence matters, and it includes examining not just sequence and structure but organization, function and expression patterns of gene also. Moreover, as Levine et al. (2006) care to admit, it is not always possible to determine whether the non-coding paralogous sequence is ancestral or the coding sequence itself. Cai et al. (2008) saliently point out that the paralogous non-coding sequence can be inferred, at certain times, as being the degenerated version of the de novo protein-coding gene.

Another issue has been supposing that any duplicated genes should be proximate to each other on the same chromosome and thereby assuming a conserved synteny has been a general approach taken when searching for potential homologs, however divergent, to candidate orphan genes. But the phenomenon of non-homologous crossing over, as well as frequent transposon-mediated relocation and chromosomal translocation, overturns this since paralogs can easily be moved even if created by tandem duplication. There may also not be an appreciation of what information non coding DNA contains. Fven though the ENCODE consortium (2012) essentially settled the fact that large swathes of the non-coding genome are probably functional, there remains a dogma that intergenic (and intronic) DNA fundamentally consists of randomly arranged, functionless and basically unconserved sequences. Weisman (2022) repeatedly refers to protein-coding de novo genes as being essentially “random” amino acid sequences and compares them to those in random sequence libraries used in experiments. Bornberg-Bauer et al. (2021) also claim that de novo proteins are indistinguishable from those of randomly obtained sequences. Whilst it is true that at least a significant portion of ncDNA may just be “spacer sequence”, not conserved except perhaps for length, most genomes are replete with vast numbers of functional non-coding elements that include binding sites for transcription factors, RNA transcripts involved in regulating gene expression, and of course, many pseudogenes.

This limited study, of course, did not have the opportunity to examine claims made about all purportedly de novo genes. Only an exhaustive meta-analysis could possibly determine this. Moreover, the results presented here are also contestable, especially given the lack of substantial statistical evidence in places as well as experimental confirmation. But the criterion for homology detection used here has been plausibility, not certainty: all four de novo genes examined here have credible homologs that could have served as evolutionary precursors. And if such homologs potentially exist, then it is incumbent on those making the claim about de novo origins to definitively rule them out since a de novo gene should not resemble any other. This, unfortunately, has not been done in nearly every reported case despite it being more parsimonious to infer that an old gene has been revived than to have emerged from non-genic DNA.

The approach used here to detect homologs can be properly criticized as “flexible” maybe lacking controls, but it is nonetheless straight forward. In recognition of the inherent limitations of sequence analysis algorithms, sub-sequences of the target gene’s products were used rather than whole ones. The N-terminus proved a sensible choice given its relative importance, but other regions could also have been included. This meant that potential homologs showed up that could have been much more divergent in other places and so liable to be filtered out altogether. The de novo genes investigated here were limited to protein-coding genes, not to ncRNA genes which may be of considerable significance. Examples of this include Poldi in *Mus* (Heinen et al. 2009); some miRNA genes are believed to have arisen de novo (Johnson et al. 2022) although (as is explained above) several studies also have reported that degenerated pseudogenes can exhibit RNA expression (Hirotsune et al. 2003; Piehler et al. 2006).

The problems associated with the birth of a new protein-coding gene from an arbitrary stretch of intergenic DNA are still quite formidable. The acquisition of transcriptional activity and presence of a viable ORF is the relatively easy part. The difficulty arises in that the coding region often requires a signal peptide for the cell to know where to transport it, and there must exist some structural motifs to allow it to fold correctly. From a biophysical perspective, the emergence of novel functional proteins from random DNA stretches is difficult to explain (Bornberg-Bauer et al 2015). Weisman (2022) highlights research showing, however, that some randomly arranged heteropolymers have these properties (Heames et al. 2022). A de novo protein-coding gene would also need 5’ and 3’ UTRs to support translational efficiency and the stability of the mRNA, along with a Kozak sequence to initiate translation (although this can easily be acquired). Even if the gene perform does some function, it does not follow that it would be useful, thereby allowing it to be preserved by natural selection. The de novo recruitment of some ncDNA into an existing protein-coding gene is entirely possible, such as through retrotransposon insertion (Poretti et al 2023; Yang et al 2021), although this describes an expansion of an already functioning gene rather than one that is born “from scratch”. Also, if a peptide is, in fact, confirmed not to exist anywhere else within the protein repertoire, this may not by itself be that significant. In a study reporting the presence of candidate de novo genes in Saccharomyces, Blevins et al. (2019) state that “the data suggest that whereas some of these (de novo) loci are likely to translate for peptides with biological significance, others may be transiently translated without ever becoming functional”. As such, far from being a source of functional diversity, contributing to evolutionary innovation and adaptation, some translated de novo transcripts may prove to be the source of frequent background noise. The peptides that they encode, which could be toxic, may float about in the cytoplasm before being tagged and sent to the proteasome for degrading.

Recently, several studies have been conducted that appear to reveal that functional microproteins may have arisen de novo from lncRNAs whose coding potential has hitherto been largely ignored (Pan et al 2022). They have also been implicated in pathogenesis (Xing et al 2022). Specifically, Vakirlis et al (2022) and An et al (2023) report experimental evidence from cell growth phenotypes and transgenic studies in mice that there are micropeptides synthesized in the human lineage alone that may contribute to human brain size and other distinctive traits. Peptide verification using ribosome profiling and immunofluorescence staining, however, can prove to be erroneous and the independent reproducibility of these experiments is questionable. Differences in the genetic background and inherent variability across cell lines can lead to a misinterpretation of phenotypes (Kim et al 2020); organs like the brain in mice and humans may be too divergent, and intraspecies variation too great, to draw conclusions about functional effects. Moreover, as Vakirlis et al (2022) acknowledge, novel ORFs may potentially emerge out of pseudogenic loci, as in the case of their exemplar transcript which is expressed in the heart and has all the hallmarks of being a pseudogene that was later re-purposed as a regulatory lncRNA. Indeed, Schmitz and Bornberg-Bauer (2017) notably admit that, “a (partial) revival of pseudogenized gene” is a possibility in molecular evolution and that certain “fragments of a pseudogenized gene that has been somewhat eroded by drift could become part of a de novo ORF.”

Whatever the truth about micropeptide origins, the claim that protein-coding genes can arise from DNA, with entirely novel ORFs, and that these were always non-coding, has not been supported with the necessary confirmatory information. Also, declaring a gene to have emerged de novo, without any homologs, has the drawback of hindering determining its functionality, assuming that it would resemble any orthologs. A final point of interest beyond the scope here concerns whether diverged pseudogenes, lying dormant or not fully expressed, could be reactivated at a later point in time, including within humans. This is especially interesting given the disputed claim that human evolution has essentially come to a halt (Templeton 2010). However, any reactivations may well affect future evolutionary trajectories especially if the revived gene has changed much in function and it is able to confer a survival advantage.

## Methods and Materials

Genomic and proteomic data was downloaded from both NCBI and Ensembl. The accession numbers for genes/proteins referred to in Figures 1 - 6 are given as follows: FLJ33706 in H. sapiens (NP_872390), C20orf203 in P. troglodytes (XR_002940896), ZNF678.1 in H. sapiens (NP_848644), Goddard in D. melanogaster (NP_648713), Protein Max in D. melanogaster (NP_001287114), Protein Max in C. capitata (JAC03861), Protein MAX in M. domestica (XP_005180678), BSC4 in S. cerevisiae (ACB40946), Vps38 in S. cerevisiae (QHB10501), Vps38 in S. paradoxus (XP_033768095), AFGP2 in G. morhua (QBH74515) AFGP2 in B. saida (QBH74504), APOA2 in G. morhua (XP_030236468) and APOA1 in D. rerio (NP_571203). Other genes cited for a dubious status are CLLU1 (NR_027932) and C22orf45 (NR_028484). Images used in the figures for the predicted proteins of Goddard and Protein Max in D. melanogaster were borrowed from Uniprot (Q9VUG4) and (P91664), respectively.

Sequence analysis and motif search was performed with Interpro (Mcdowall and Hunter 2003) and the Conserved Domains Database (CDD) (Marchler-Bauer et al. 2015) (https://www.ncbi.nlm.nih.gov/cdd/). Homology detection was conducted with BLAST (Altschul et al. 1997), and also BLASTN, using the following adjusted parameters: BLOSUM60 matrix; word size =3; Gap penalty 11 and extension of 2. As stated above, the N-termini of the candidate de novo genes were used when searching. In the case of FLJ33706, the whole sequence was used for BLASTP and PSI-BLAST for up to 5 iterations. Amino acid composition was determined using Protparam (https://web.expasy.org/protparam/) which is part of the ExPASy server’s suite of tools (Gasteiger et al. 2003). Nucleotide alignments were made using the T-COFFEE (Di Tomasso et al. 2011) server (http://www.tcoffee.crg.cat/). Codon usage analysis was performed using the Sequence Manipulation Suite (Stothard 2000) (https://www.bioinformatics.org/sms2/codon_usage.html) Pairwise amino acid alignments were made using the SIM tool (https://web.expasy.org/sim/) also from the ExPASy server. BLOSUM30 was used as the substitution matrix because it is the most suitable one available for any two sequences that are expected to show the lowest sequence identity (Baussand and Carbone 2008), typically evolutionarily distant homologs sharing a remote ancestry or rapidly diverged paralogs. The gap opening and extension penalties were both set to 2, allowing for the presence of significant indels. Multiple alignments, both nucleotide and amino acid, were created using the CLUSTAL algorithm (Sievers et al. 2011) (https://www.ebi.ac.uk/Tools/msa/clustalo/). Primary sequence alignments, as shown in the figures, were visualized with Word Art.

Statistical prediction of tertiary structure used Phyre2 (Kelley et al. 2015) (http://www.sbg.bio.ic.ac.uk/phyre2/html/page.cgi?id=index), availing of the normal modelling mode of HMMs for fold recognition. In the case of Goddard and MAX in Drosophila, Alphafold/Uniprot entries for pdb files (Q9VUG4 and M9NDC9) were used instead due to higher quality. AFGP2 in the Atlantic cod (A0A481T0Z2) was also taken from Alphafold. All Pdb files used are provided as a zip file. Visualization of the structural data was performed using SWISS-MODEL (Waterhouse et al 2018) (https://swissmodel.expasy.org/). Structural alignment, including the determination of the RMSD, was conducted with the TM-align package (Zhang et al 2005) (https://zhanggroup.org/TM-align/) and 3D-Match from the Softberry suite (http://www.softberry.com/berry.phtml?topic=3dmatch&group=help&subgroup=3d-expl). Secondary protein structure was predicted using PSI-PRED (McGuffin et al 2000) (http://bioinf.cs.ucl.ac.uk/psipred/). Levels of protein aggregation and disorder were calculated using PASTA 2.0 (Walsh et al 2014) (http://old.protein.bio.unipd.it/pasta2) and flDPnn (Hu et al 2021) http://biomine.cs.vcu.edu/servers/flDPnn/. Lipid-binding sites were predicted with DisoLipPred (Katuwawala et al 2021) (http://biomine.cs.vcu.edu/servers/DisoLipPred). Predictions of DNA-binding sites used DNAGenie (Zhang et al 2021) (http://biomine.cs.vcu.edu/servers/DNAgenie). RNA-binding status was predicted with RNAPred (Kumar et al 2010) (https://webs.iiitd.edu.in/raghava/rnapred) and DRNAPred (Yan et al 2017) (http://biomine.cs.vcu.edu/servers/DRNApred/). In the case of the former, default settings were used for the composition with a higher sensitivity setting for the PSSM.

Subcellular localization prediction availed of Wolf PSort (Horton et al 2007) (https://www.genscript.com/wolf-psort.html) and TargetP-2.0 (Armenteros et al 2019) (https://services.healthtech.dtu.dk/services/TargetP-2.0/). Mitochondrial transfer peptide prediction was performed using DeepMito (Savojardo et al 2019) (http://busca.biocomp.unibo.it/deepmito/) and MitPed (Kumar et al 2005) (https://webs.iiitd.edu.in/raghava/mitpred/submit.html). Nuclear localization signal prediction was implemented with the NLStradamus server (Nguyen Ba at al 2009) (http://www.moseslab.csb.utoronto.ca/NLStradamus) using only default parameters. Prediction of function used FFPRED3 (http://bioinf.cs.ucl.ac.uk/psipred) (Cozzetto et al 2016) and transcription factor identity was classified with DeepTFactor (Kim et al 2020), a deep learning neural network. P-values for the significance of the amino acid composition of proteins, relative to predictions in the genetic code, were calculated using a standard Pearson chi square test explicated by Bozorgmehr (2015). Correlation coefficients for the amino acid composition comparison used Pearson as the method.

## Acknowledgements

This study received no funding or support. I am, however, grateful for many relevant comments shared directly with me by Dr. Caroline Weisman of Princeton University. I would also like to thank Prof. Ralph Bundschuh of Ohio State University for allowing me to reproduce mathematical equations related to the use of BLAST and PSI-BLAST.

## Statements and Declarations

The sole author has no relevant financial or non-financial competing interests to disclose and did not receive funding from any organization for conducting this study.

## References

1. Ajazi A, Bruhn C, Shubassi G, Lucca C, Ferrari E, Cattaneo A, Bachi A, Manfrini N, Biffo S, Martini E, Minucci S, Vernieri C, Foiani M. Endosomal trafficking and DNA damage checkpoint kinases dictate survival to replication stress by regulating amino acid uptake and protein synthesis. Dev Cell. 2021 Sep 27;56(18):2607–2622.e6. doi: 10.1016/j.devcel.2021.08.019. Epub 2021 Sep 16. PMID: 34534458.

2. Alawad A, Alharbi S, Alhazzaa O, Alagrafi F, Alkhrayef M, Alhamdan Z, Alenazi A, Al-Johi H, Alanazi IO, Hammad M. Phylogenetic and Structural Analysis of the Pluripotency Factor Sex-Determining Region Y box2 Gene of Camelus dromedarius (cSox2). Bioinform Biol Insights. 2016 Jul 26;10:111–20. doi: 10.4137/BBI.S39047.

3. Altschul SF, Madden TL, Schäffer AA, Zhang J, Zhang Z, Miller W, Lipman DJ. Gapped BLAST and PSI-BLAST: a new generation of protein database search programs. Nucleic Acids Res. 1997 Sep 1;25(17):3389–402. doi: 10.1093/nar/25.17.3389.

4. An NA, Zhang J, Mo F, Luan X, Tian L, Shen QS, Li X, Li C, Zhou F, Zhang B, Ji M, Qi J, Zhou WZ, Ding W, Chen JY, Yu J, Zhang L, Shu S, Hu B, Li CY. De novo genes with an lncRNA origin encode unique human brain developmental functionality. Nat Ecol Evol. 2023 Feb;7(2):264–278. doi: 10.1038/s41559-022-01925-6. Epub 2023 Jan 2.

5. Armenteros JJ, Salvatore M, Emanuelsson O, Winther O, von Heijne G, Elofsson A, Nielsen H. Detecting sequence signals in targeting peptides using deep learning. Life Sci Alliance. 2019 Sep 30;2(5):e201900429. doi: 10.26508/lsa.201900429.

6. Assis R, Kondrashov AS, Koonin EV, Kondrashov FA. Nested genes and increasing organizational complexity of metazoan genomes. Trends Genet. 2008 Oct;24(10):475–8. doi: 10.1016/j.tig.2008.08.003. Epub 2008 Sep 5.

7. Assis R, Bachtrog D. Neofunctionalization of young duplicate genes in Drosophila. Proc Natl Acad Sci U S A. 2013 Oct 22;110(43):17409–14. doi: 10.1073/pnas.1313759110. Epub 2013 Oct 7. PMID: 24101476; PMCID: PMC3808614.

8. Baalsrud HT, Tørresen OK, Solbakken MH, Salzburger W, Hanel R, Jakobsen KS, Jentoft S. De Novo Gene Evolution of Antifreeze Glycoproteins in Codfishes Revealed by Whole Genome Sequence Data. Mol Biol Evol. 2018 Mar 1;35(3):593–606. doi: 10.1093/molbev/msx311.

9. Bai Y, Casola C, Betrán E. Evolutionary origin of regulatory regions of retrogenes in Drosophila. BMC Genomics. 2008 May 22;9:241. doi: 10.1186/1471-2164-9-241.

10. Barthez M, Poplineau M, Elrefaey M, Caruso N, Graba Y, Saurin AJ. Human ZKSCAN3 and Drosophila M1BP are functionally homologous transcription factors in autophagy regulation. Sci Rep. 2020 Jun 15;10(1):9653. doi: 10.1038/s41598-020-66377-z.

11. Basile W, Salvatore M, Elofsson A. The classification of orphans is improved by combining searches in both proteomes and genomes. BioRxiv. 2019 doi: 10.1101/185983

12. Bazykin GA, Kochetov AV. Alternative translation start sites are conserved in eukaryotic genomes. Nucleic Acids Res. 2011 Jan;39(2):567–77. doi: 10.1093/nar/gkq806. Epub 2010 Sep 22.

13. Baussand J, Carbone A. Inconsistent distances in substitution matrices can be avoided by properly handling hydrophobic residues. Evol Bioinform Online. 2008 Oct 9;4:255–61. doi: 10.4137/ebo.s885.

14. Blevins WR, Ruiz-Orera J, Messeguer X, et al. Frequent birth of de novo genes in the compact yeast genome. bioRxiv; 2019. DOI: 10.1101/575837.

15. Blevins WR, Ruiz-Orera J, Messeguer X, Blasco-Moreno B, Villanueva-Cañas JL, Espinar L, Díez J, Carey LB, Albà MM. Uncovering de novo gene birth in yeast using deep transcriptomics. Nat Commun. 2021 Jan 27;12(1):604. doi: 10.1038/s41467-021-20911-3. PMID: 33504782; PMCID: PMC7841160.

16. Boratyn GM, Schäffer AA, Agarwala R, Altschul SF, Lipman DJ, Madden TL. Domain enhanced lookup time accelerated BLAST. Biol Direct. 2012 Apr 17;7:12. doi: 10.1186/1745-6150-7-12. PMID: 22510480; PMCID: PMC3438057.

17. Bornberg-Bauer E, Hlouchova K, Lange A. Structure and function of naturally evolved de novo proteins. Curr Opin Struct Biol. 2021 8:175–183. doi: 10.1016/j.sbi.2020.11.010.

18. Bornberg-Bauer E, Schmitz J, Heberlein M. Emergence of de novo proteins from ’dark genomic matter’ by ’grow slow and moult’. Biochem Soc Trans. 2015 Oct;43(5):867–73. doi: 10.1042/BST20150089.

19. Bozorgmehr JH. The origin of chromosomal histones in a 30S ribosomal protein. Gene. 2019 Feb 5;726:144155. doi: 10.1016/j.gene.2019.144155.

20. Bozorgmehr JH. Quantifying protein sequences with reference to the genetic code. J Theor Biol. 2015 May 7;372:39–46. doi: 10.1016/j.jtbi.2015.02.017.

21. Broeils LA, Ruiz-Orera J, Snel B, Hubner N, van Heesch S. Evolution and implications of de novo genes in humans. Nat Ecol Evol. 2023. doi: 10.1038/s41559-023-02014-y.

22. Brosch M, Saunders GI, Frankish A, Collins MO, Yu L, Wright J, Verstraten R, Adams DJ, Harrow J, Choudhary JS, Hubbard T (2011). Shotgun proteomics aids discovery of novel protein-coding genes, alternative splicing, and “resurrected” pseudogenes in the mouse genome. Genome Res;21(5):756–67. doi: 10.1101/gr.114272.110.

23. Bungard D et al (2017) Foldability of a natural de novo evolved protein. Structure 25(11):1687–1696

24. Bustamante CD, Nielsen R, Hartl DL. A maximum likelihood method for analyzing pseudogene evolution: implications for silent site evolution in humans and rodents. Mol Biol Evol. 2002 Jan;19(1):110–7. doi: 10.1093/oxfordjournals.molbev.a003975.

25. Cai J, Zhao R, Jiang H,Wang W (2008) De novo origination of a new protein-coding gene in Saccharomyces cerevisiae. Genetics 179, 487–496

26. Carelli FN, Hayakawa T, Go Y, Imai H, Warnefors M, Kaessmann H. The life history of retrocopies illuminates the evolution of new mammalian genes. Genome Res. 2016 Mar;26(3):301–14. doi: 10.1101/gr.198473.115.

27. Casci T. A gene is born. Nat Rev Genet 9, 415 (2008). 10.1038/nrg2394

28. Casola C. From De Novo to “De Nono”: The Majority of Novel Protein-Coding Genes Identified with Phylostratigraphy Are Old Genes or Recent Duplicates. Genome Biol Evol. 2018 Nov 1;10(11):2906–2918. doi: 10.1093/gbe/evy231

29. Cassiday LA, Maher LJ 3rd. Having it both ways: transcription factors that bind DNA and RNA. Nucleic Acids Res. 2002 Oct 1;30(19):4118–26. doi: 10.1093/nar/gkf512.

30. Carvunis A-R et al (2012) Proto-genes and de novo gene birth. Nature 487(7407):370

31. Chandrasekar V, Dreyer JL. The Brain-Specific Neural Zinc Finger Transcription Factor 2b (NZF-2b/7ZFMyt1) Suppresses Cocaine Self-Administration in Rats. Front Behav Neurosci. 2010 Apr 5;4:14. doi: 10.3389/fnbeh.2010.00014. PMID: 20407577;

32. Chen L, DeVries AL, Cheng C-HC (1997) Evolution of antifreeze glycoprotein gene from a trypsinogen gene in Antarctic notothenioid fish. Proc Natl Acad Sci 94(8):3811–3816

33. Cheng C-HC (1998) Evolution of the diverse antifreeze proteins. Curr Opin Genet Dev 8(6):715–720

34. Cherezov, R.O., Vorontsova, J.E. & Simonova, O.B. The Phenomenon of Evolutionary “*De Novo* Generation” of Genes. Russ J Dev Biol 52, 390–400 (2021). 10.1134/S1062360421060035

35. Choo KH, Tan TW, Ranganathan S. A comprehensive assessment of N-terminal signal peptides prediction methods. BMC Bioinformatics. 2009 Dec 3;10 Suppl 15(Suppl 15):S2. doi: 10.1186/1471-2105-10-S15-S2. PMID: 19958512; PMCID: PMC2788353.

36. Ciomborowska J, Rosikiewicz W, Szklarczyk D, Makałowski W, Makałowska I. “Orphan” retrogenes in the human genome. Mol Biol Evol. 2013 Feb;30(2):384–96. doi: 10.1093/molbev/mss235. Epub 2012 Oct 12. PMID: 23066043; PMCID: PMC3548309.

37. Comas D, Plaza S, Calafell F, Sajantila A, Bertranpetit J. Recent insertion of an Alu element within a polymorphic human-specific Alu insertion. Mol Biol Evol. 2001 Jan;18(1):85–8. doi: 10.1093/oxfordjournals.molbev.a003722. PMID: 11141195.

38. Cozzetto D, Minneci F, Currant H, Jones DT. FFPred 3: feature-based function prediction for all Gene Ontology domains. Sci Rep. 2016 Aug; 6:31865. doi: 10.1038/srep31865.

39. Deng C, Cheng CH, Ye H, He X, Chen L. Evolution of an antifreeze protein by neofunctionalization under escape from adaptive conflict. Proc Natl Acad Sci U S A. 2010 Dec 14;107(50):21593–8. doi: 10.1073/pnas.1007883107.

40. Deng G, Andrews DW, Laursen RA. Amino acid sequence of a new type of antifreeze protein, from the longhorn sculpin Myoxocephalus octodecimspinosis. FEBS Lett. 1997 Jan 27;402(1):17–20. doi: 10.1016/s0014-5793(96)01466-4. PMID: 9013849.

41. Desiere F, Deutsch EW, King NL, Nesvizhskii AI, Mallick P, Eng J, Chen S, Eddes J, Loevenich SN, Aebersold R. The PeptideAtlas project. Nucleic Acids Res. 2006 Jan 1;34(Database issue): D655–8. doi: 10.1093/nar/gkj040.

42. Ding F, Borreguero JM, Buldyrey SV, Stanley HE, Dokholyan NV. Mechanism for the alpha-helix to beta-hairpin transition. Proteins. 2003 Nov 1;53(2):220–8. doi: 10.1002/prot.10468. PMID: 14517973.

43. Di Tommaso P, Moretti S, Xenarios I, Orobitg M, Montanyola A, Chang JM, Taly JF, Notredame C. T-Coffee: a web server for the multiple sequence alignment of protein and RNA sequences using structural information and homology extension. Nucleic Acids Res. 2011 Jul;39(Web Server issue): W13–7. doi: 10.1093/nar/gkr245.

44. Dotiwala F, Eapen VV, Harrison JC, Arbel-Eden A, Ranade V, Yoshida S, Haber JE. DNA damage checkpoint triggers autophagy to regulate the initiation of anaphase. Proc Natl Acad Sci U S A. 2013 Jan 2;110(1): E41–9. doi: 10.1073/pnas.1218065109.

45. Dyson HJ, Wright PE. Elucidation of the protein folding landscape by NMR. Methods Enzymol. 2005; 394:299–321. doi: 10.1016/S0076-6879(05)94011-1. PMID: 15808225.

46. Eliopoulos AG, Havaki S, Gorgoulis VG. DNA Damage Response and Autophagy: A Meaningful Partnership. Front Genet. 2016: 21;7:204. doi: 10.3389/fgene.2016.00204.

47. ENCODE Project Consortium. An integrated encyclopedia of DNA elements in the human genome. Nature. 2012 Sep 6;489(7414):57–74. doi: 10.1038/nature11247.

48. Esfeld K, Berardi AE, Moser M, Bossolini E, Freitas L, Kuhlemeier C. Pseudogenization and Resurrection of a Speciation Gene. Curr Biol. 2018 Dec 3;28(23):3776–3786.e7. doi: 10.1016/j.cub.2018.10.019. Epub 2018 Nov 21. PMID: 30472000.

49. Farmiloe G, Lodewijk GA, Robben SF, van Bree EJ, Jacobs FMJ. Widespread correlation of KRAB zinc finger protein binding with brain-developmental gene expression patterns. Philos Trans R Soc Lond B Biol Sci. 2020 Mar 30;375(1795):20190333. doi: 10.1098/rstb.2019.0333. Epub 2020 Feb 10. PMID: 32075554; PMCID: PMC7061980.

50. Fiddes IT, Lodewijk GA, Mooring M, Bosworth CM, Ewing AD, Mantalas GL, Novak AM, van den Bout A, Bishara A, Rosenkrantz JL, Lorig-Roach R, Field AR, Haeussler M, Russo L, Bhaduri A, Nowakowski TJ, Pollen AA, Dougherty ML, Nuttle X, Addor MC, Zwolinski S, Katzman S, Kriegstein A, Eichler EE, Salama SR, Jacobs FMJ, Haussler D. Human Specific NOTCH2NL Genes Affect Notch Signaling and Cortical Neurogenesis. Cell. 2018 May 31;173(6):1356–1369.e22. doi: 10.1016/j.cell.2018.03.051.

51. Florio M, Albert M, Taverna E, Namba T, Brandl H, Lewitus E, Haffner C, Sykes A, Wong FK, Peters J, Guhr E, Klemroth S, Prüfer K, Kelso J, Naumann R, Nüsslein I, Dahl A, Lachmann R, Pääbo S, Huttner WB. Human-specific gene ARHGAP11B promotes basal progenitor amplification and neocortex expansion. Science. 2015 Mar 27;347(6229):1465–70. doi: 10.1126/science.aaa1975.

52. Fraïsse C, Puixeu Sala G, Vicoso B. Pleiotropy Modulates the Efficacy of Selection in Drosophila melanogaster. Mol Biol Evol. 2019 Mar 1;36(3):500–515. doi: 10.1093/molbev/msy246. PMID: 30590559; PMCID: PMC6389323.

53. Gauthier SY, Scotter AJ, Lin FH, Baardsnes J, Fletcher GL, Davies PL. A re-evaluation of the role of type IV antifreeze protein. Cryobiology. 2008 Dec;57(3):292–6. doi: 10.1016/j.cryobiol.2008.10.122.

54. Ghalamara, S., Silva, S., Brazinha, C. et al. Structural diversity of marine anti-freezing proteins, properties and potential applications: a review. Bioresour. Bioprocess. 9, 5 (2022). 10.1186/s40643-022-00494-7

55. Goonesekere NC, Lee B. Frequency of gaps observed in a structurally aligned protein pair database suggests a simple gap penalty function. Nucleic Acids Res. 2004 May 20;32(9):2838–43. doi: 10.1093/nar/gkh610. PMID: 15155852; PMCID: PMC419611.

56. Gotea V, Petrykowska HM, Elnitski L. Bidirectional promoters as important drivers for the emergence of species-specific transcripts. PLoS One. 2013;8(2):e57323. doi: 10.1371/journal.pone.0057323.

57. Grandchamp A, Berk K, Dohmen E, Bornberg-Bauer E. New Genomic Signals Underlying the Emergence of Human Proto-Genes. Genes (Basel). 2022 Jan 31;13(2):284. doi: 10.3390/genes13020284. PMID: 35205330; PMCID: PMC8871994.

58. Gubala AM, Schmitz JF, Kearns MJ, Vinh TT, Bornberg-Bauer E, Wolfner MF, Findlay GD. The Goddard and Saturn Genes Are Essential for Drosophila Male Fertility and May Have Arisen De Novo. Mol Biol Evol. 2017 May 1;34(5):1066–1082. doi: 10.1093/molbev/msx057. PMID: 28104747; PMCID: PMC5400382.

59. Guerzoni D, McLysaght A (2015) New genes from non-coding sequence: the role of de novo protein-coding genes in eukaryotic evolutionary innovation. Phil. Trans. R. Soc. B 370, 20140332

60. Guschanski K, Warnefors M, Kaessmann H. The evolution of duplicate gene expression in mammalian organs. Genome Res. 2017 Sep;27(9):1461–1474. doi: 10.1101/gr.215566.116. Epub 2017 Jul 25. PMID: 28743766; PMCID: PMC5580707.

61. Hangauer MJ, Vaughn IW, McManus MT. Pervasive transcription of the human genome produces thousands of previously unidentified long intergenic noncoding RNAs. PLoS Genet. 2013 Jun;9(6):e1003569. doi: 10.1371/journal.pgen.1003569.

62. Harrington JM, Nishanova T, Pena SR, Hess M, Scelsi CL, Widener J, Hajduk SL. A retained secretory signal peptide mediates high density lipoprotein (HDL) assembly and function of haptoglobin-related protein. J Biol Chem. 2014 Sep 5;289(36):24811–20. doi: 10.1074/jbc.M114.567578. Epub 2014 Jul 17.

63. Hartl DL, Taubes CH. Compensatory nearly neutral mutations: selection without adaptation. J Theor Biol. 1996 Oct 7;182(3):303–9. doi: 10.1006/jtbi.1996.0168.

64. Hartford CCR, Lal A. When Long Noncoding Becomes Protein Coding. Mol Cell Biol. 2020 Feb 27;40(6):e00528–19. doi: 10.1128/MCB.00528-19.

65. Heames B, Buchel F, Aubel M, Tretyachenko V, Loginov D, Novák P, Lange A, Bornberg-Bauer E, Hlouchová K. Experimental characterization of de novo proteins and their unevolved random-sequence counterparts. Nat Ecol Evol. 2023 Apr;7(4):570–580. doi: 10.1038/s41559-023-02010-2. Epub 2023 Apr 6.

66. Heinen TJ et al (2009) Emergence of a new gene from an intergenic region. Curr Biol 19(18):1527–1531

67. Hirotsune S, Yoshida N, Chen A, Garrett L, Sugiyama F, Takahashi S, Yagami K, Wynshaw-Boris A, Yoshiki A. An expressed pseudogene regulates the messenger-RNA stability of its homologous coding gene. Nature. 2003 May 1;423(6935):91–6. doi: 10.1038/nature01535.

68. Holmes ZE, Hamilton DJ, Hwang T, Parsonnet NV, Rinn JL, Wuttke DS, Batey RT. The Sox2 transcription factor binds RNA. Nat Commun. 2020;11(1):1805. doi: 10.1038/s41467-020-15571-8. PMID: 32286318; PMCID: PMC7156710.

69. Horton P, Park KJ, Obayashi T, Fujita N, Harada H, Adams-Collier CJ, Nakai K. WoLF PSORT: protein localization predictor. Nucleic Acids Res. 2007 Jul;35(Web Server issue):W585–7. doi: 10.1093/nar/gkm259.

70. Hu G, Katuwawala A, Wang K, Wu Z, Ghadermarzi S, Gao J, Kurgan L. flDPnn: Accurate intrinsic disorder prediction with putative propensities of disorder functions. Nat Commun. 2021 Jul 21;12(1):4438. doi: 10.1038/s41467-021-24773-7.

71. Jacob F (1977) Evolution and tinkering. Science 196(4295):1161–1166

72. Ji Z, Song R, Regev A, Struhl K (2015) Many lncRNAs, 5′UTRs, and pseudogenes are translated and some are likely to express functional proteins. eLife 4, e08890

73. Kang LF, Zhu ZL, Zhao Q, Chen LY, Zhang Z. Newly evolved introns in human retrogenes provide novel insights into their evolutionary roles. BMC Evol Biol. 2012 Jul 28;12:128. doi: 10.1186/1471-2148-12-128.

74. Kato GJ, Lee WM, Chen LL, Dang CV. Max: functional domains and interaction with c Myc. Genes Dev. 1992 Jan;6(1):81–92. doi: 10.1101/gad.6.1.81. PMID: 1730412.

75. Kast DJ, Dominguez R. The Cytoskeleton-Autophagy Connection. Curr Biol. 2017 Apr 24;27(8):R318–R326. doi: 10.1016/j.cub.2017.02.061.

76. Kelley LA, Mezulis S, Yates CM, Wass MN, Sternberg MJ. The Phyre2 web portal for protein modeling, prediction and analysis. Nat Protoc. 2015 Jun;10(6):845–58. doi: 10.1038/nprot.2015.053. Epub 2015 May 7. PMID: 25950237; PMCID: PMC5298202.

77. Kaessmann H. Origins, evolution, and phenotypic impact of new genes. Genome Res. 2010 Oct;20(10):1313–26. doi: 10.1101/gr.101386.109. Epub 2010 Jul 22

78. Katuwawala A, Zhao B, Kurgan L. DisoLipPred: accurate prediction of disordered lipid binding residues in protein sequences with deep recurrent networks and transfer learning. Bioinformatics. 2021;38(1):115–124. doi: 10.1093/bioinformatics/btab640.

79. Khan C, Muliyil S, Rao BJ. Genome Damage Sensing Leads to Tissue Homeostasis in Drosophila. Int Rev Cell Mol Biol. 2019; 345:173–224. doi: 10.1016/bs.ircmb.2018.12.001. Epub 2019 Jan 14. PMID: 30904193.

80. Khor BY, Tye GJ, Lim TS, Choong YS. General overview on structure prediction of twilight-zone proteins. Theor Biol Med Model. 2015 Sep 4; 12:15. doi: 10.1186/s12976-015-0014-1.

81. Kim GB, Gao Y, Palsson BO, Lee SY. DeepTFactor: A deep learning-based tool for the prediction of transcription factors. Proc Natl Acad Sci U S A. 2021 Jan 12;118(2):e2021171118. doi: 10.1073/pnas.2021171118.

82. Kimura M. (1964). Diffusion models in population genetics. Journal of Applied Probability, 1(2), 177–232. doi:10.2307/3211856

83. Kim J, Koo BK, Knoblich JA. Human organoids: model systems for human biology and medicine. Nat Rev Mol Cell Biol. 2020 Oct;21(10):571–584. doi: 10.1038/s41580-020-0259-3. Epub 2020 Jul 7. PMID: 32636524; PMCID: PMC7339799.

84. Kiselak EA, Shen X, Song J, Gude DR, Wang J, Brody SL, Strauss JF 3rd, Zhang Z. Transcriptional regulation of an axonemal central apparatus gene, sperm-associated antigen 6, by a SRY-related high mobility group transcription factor, S-SOX5. J Biol Chem. 2010 Oct 1;285(40):30496–505. doi: 10.1074/jbc.M110.121590.

85. Kowalczyk MS, Hughes JR, Garrick D, et al. Intragenic enhancers act as alternative promoters. Mol Cell. 2012;45(4):447–458. doi:10.1016/j.molcel.2011.12.021

86. Kramer MH, Farré JC, Mitra K, Yu MK, Ono K, Demchak B, Licon K, Flagg M, Balakrishnan R, Cherry JM, Subramani S, Ideker T. Active Interaction Mapping Reveals the Hierarchical Organization of Autophagy. Mol Cell. 2017 Feb 16;65(4):761–774.e5. doi: 10.1016/j.molcel.2016.12.024.

87. Hunter R. Johnson, Jessica A. Blandino, Beatriz C. Mercado, José A. Galván, William J. Higgins, Nathan H. Lents. 2022. The evolution of de novo human-specific microRNA genes on chromosome 21. American Journal of Biological Anthropology

88. Johnson BR. Taxonomically Restricted Genes Are Fundamental to Biology and Evolution. Front Genet. 2018; 9:407. doi: 10.3389/fgene.2018.00407.

89. Krieger F, Möglich A, Kiefhaber T. Effect of proline and glycine residues on dynamics and barriers of loop formation in polypeptide chains. J Am Chem Soc. 2005 Mar 16;127(10):3346–52. doi: 10.1021/ja042798i. PMID: 15755151.

90. Knowles DG, McLysaght, A (2009) Recent de novo origin of human protein-coding genes. Genome Res. 19, 1752–1759

91. Kufareva I, Abagyan R. Methods of protein structure comparison. Methods Mol Biol. 2012;857:231–57. doi: 10.1007/978-1-61779-588-6_10.

92. Kumar M, Verma R, Raghava GP. Prediction of mitochondrial proteins using support vector machine and hidden Markov model. J Biol Chem. 2006 Mar 3;281(9):5357-63. doi: 10.1074/jbc.M511061200. Epub 2005 Dec 8. PMID: 16339140.

93. Kumar A. An overview of nested genes in eukaryotic genomes. Eukaryot Cell. 2009 Sep;8(9):1321–9. doi: 10.1128/EC.00143-09.

94. Kumar M, Gromiha MM, Raghava GP. SVM based prediction of RNA-binding proteins using binding residues and evolutionary information. J Mol Recognit. 2011 Mar-Apr;24(2):303–13. doi: 10.1002/jmr.1061.

95. Lange A, Patel PH, Heames B, Damry AM, Saenger T, Jackson CJ, Findlay GD, Bornberg-Bauer E. Structural and functional characterization of a putative de novo gene in Drosophila. Nat Commun. 2021;12(1):1667. doi: 10.1038/s41467-021-21667-6.

96. Lee MM, Chan MK, Bundschuh R. Simple is beautiful: a straightforward approach to improve the delineation of true and false positives in PSI-BLAST searches. Bioinformatics. 2008 Jun 1;24(11):1339–43. doi: 10.1093/bioinformatics/btn130.

97. Lee YC, Chang HH. The evolution and functional significance of nested gene structures in Drosophila melanogaster. Genome Biol Evol. 2013;5(10):1978–85. doi: 10.1093/gbe/evt149.2188. PMID: 24084778; PMCID: PMC3814207.

98. Lee AM, Wu CT. Enhancer-promoter communication at the yellow gene of Drosophila melanogaster: diverse promoters participate in and regulate trans interactions. Genetics. 2006 Dec;174(4):1867–80. doi: 10.1534/genetics.106.064121.

99. Levine MT, Jones CD, Kern AD, Lindfors HA, Begun DJ. Novel genes derived from noncoding DNA in Drosophila melanogaster are frequently X-linked and exhibit testis biased expression. Proc Natl Acad Sci USA. 2006;103(26):9935–9. doi: 10.1073/pnas.0509809103.

100. Li WH, Gojobori T, Nei M. Pseudogenes as a paradigm of neutral evolution. Nature. 1981 Jul 16;292(5820):237–9. doi: 10.1038/292237a0. PMID: 7254315.

101. Li CY, ZhangY, Wang Z, ZhangY, Cao C, Zhang PW, Lu SJ, Li XM, Yu Q, Zheng X, Du Q, Uhl GR, Liu QR. Wei LA (2010) Human-specific de novo protein-coding gene associated with human brain functions. PLoS Comput Biol 26;6(3)

102. Li C, Zhang J. Stop-codon read-through arises largely from molecular errors and is generally nonadaptive. PLoS Genet. 2019 May 23;15(5):e1008141. doi: 10.1371/journal.pgen.1008141. PMID: 31120886; PMCID: PMC6550407.

103. Liu F, Hu W, Vierstra RD. The Vacuolar Protein Sorting-38 Subunit of the *Arabidopsis* Phosphatidylinositol-3-Kinase Complex Plays Critical Roles in Autophagy, Endosome Sorting, and Gravitropism. Front Plant Sci. 2018;9:781. doi: 10.3389/fpls.2018.00781.

104. Madej T, Panchenko AR, Chen J, Bryant SH. Protein homologous cores and loops: important clues to evolutionary relationships between structurally similar proteins. BMC Struct Biol. 2007 Apr 10;7:23. doi: 10.1186/1472-6807-7-23. PMID: 17425794;

105. Madej T, Lanczycki CJ, Zhang D, Thiessen PA, Geer RC, Marchler-Bauer A, Bryant SH. MMDB and VAST+: tracking structural similarities between macromolecular complexes. Nucleic Acids Res. 2014 Jan;42(Database issue):D297–303. doi: 10.1093/nar/gkt1208. Dec 6. PMID: 24319143; PMCID: PMC3965051.

106. Mao MG, Chen Y, Liu RT, Lü HQ, Gu J, Jiang ZQ, Jiang JL. Transcriptome from Pacific cod liver reveals types of apolipoproteins and expression analysis of AFP-IV, structural analogue with mammalian ApoA-I. Comp Biochem Physiol Part D Genomics Proteomics. 2018 Dec;28:204–212. doi: 10.1016/j.cbd.2018.10.001.

107. Marchler-Bauer A, Derbyshire MK, Gonzales NR, Lu S, Chitsaz F, Geer LY, Geer RC, He J, Gwadz M, Hurwitz DI, Lanczycki CJ, Lu F, Marchler GH, Song JS, Thanki N, Wang Z, Yamashita RA, Zhang D, Zheng C, Bryant SH. CDD: NCBI’s conserved domain database. Nucleic Acids Res. 2015 Jan;43(Database issue):D222–6. doi: 10.1093/nar/gku1221.

108. Margolin JF, Friedman JR, Meyer WK, Vissing H, Thiesen HJ, Rauscher FJ 3rd. Krüppel associated boxes are potent transcriptional repression domains. Proc Natl Acad Sci U S A. 1994 May 10;91(10):4509–13. doi: 10.1073/pnas.91.10.4509.

109. Marsch-Martínez N, Reyes-Olalde JI, Chalfun-Junior A, Bemer M, Durán-Medina Y, Ochoa-Sánchez JC, Guerrero-Largo H, Herrera-Ubaldo H, Mes J, Chacón A, Escobar-Guzmán R, Pereira A, Herrera-Estrella L, Angenent GC, Delaye L, de Folter S. Twisting development, the birth of a potential new gene. iScience. 2022 Nov 19;25(12):105627. doi: 10.1016/j.isci.2022.105627. PMID: 36465114; PMCID: PMC9713375.

110. McDowall J, Hunter S. InterPro protein classification. Methods Mol Biol. 2011; 694:37–47. doi: 10.1007/978-1-60761-977-2_3. PMID: 21082426.

111. McLean P, BLAST. https://www.ndsu.edu/pubweb/~mcclean/plsc411/Blast-explanation-lecture-and-overhead.pdf

112. McGuffin LJ, Bryson K, Jones DT. The PSIPRED protein structure prediction server. Bioinformtics. 2000 Apr;16(4):404–5. doi: 10.1093/bioinformatics/16.4.404.

113. McLysaght A, Hurst LD. Open questions in the study of de novo genes: what, how and why. Nat Rev Genet. 2016 Sep;17(9):567–78. doi: 10.1038/nrg.2016.78.

114. Menlove KJ, Clement M, Crandall KA. Similarity searching using BLAST. Methods Mol Biol. 2009;537:1–22. doi: 10.1007/978-1-59745-251-9_1. PMID: 19378137.

115. Milligan MJ, Lipovich L. Pseudogene-derived lncRNAs: emerging regulators of gene expression. Front Genet. 2015 Feb 4;5:476. doi: 10.3389/fgene.2014.00476.

116. Mott R. Local sequence alignments with monotonic gap penalties. Bioinformatics. 1999 Jun;15(6):455–62. doi: 10.1093/bioinformatics/15.6.455. PMID: 10383814.

117. Moyers BA, Zhang J. Evaluating Phylostratigraphic Evidence for Widespread De Novo Gene Birth in Genome Evolution. Mol Biol Evol. 2016 May;33(5):1245–56. doi: 10.1093/molbev/msw008. Epub 2016 Jan 11. PMID: 26758516; PMCID: PMC5010002.

118. Muñoz-Gómez SA, Bilolikar G, Wideman JG, Geiler-Samerotte K. Constructive Neutral Evolution 20 Years Later. J Mol Evol. 2021 Apr;89(3):172–182. doi: 10.1007/s00239-021-09996-y. Epub 2021 Feb 19. PMID: 33604782; PMCID: PMC7982386.

119. Namy O, Duchateau-Nguyen G, Hatin I, Hermann-Le Denmat S, Termier M, Rousset JP. Identification of stop codon readthrough genes in Saccharomyces cerevisiae. Nucleic Acids Res. 2003 May 1;31(9):2289–96. doi: 10.1093/nar/gkg330.

120. Nowick K, Hamilton AT, Zhang H, Stubbs L. Rapid sequence and expression divergence suggest selection for novel function in primate-specific KRAB-ZNF genes. Mol Biol Evol. 2010 Nov;27(11):2606–17. doi: 10.1093/molbev/msq157. Epub 2010 Jun 23.

121. Nguyen Ba AN, Pogoutse A, Provart N, Moses AM. NLStradamus: a simple Hidden Markov Model for nuclear localization signal prediction. BMC Bioinformatics. 2009 Jun 29;10:202. doi: 10.1186/1471-2105-10-202. PMID: 19563654; PMCID: PMC2711084.

122. Ochoa A, Storey JD, Llinás M, Singh M. Beyond the E-Value: Stratified Statistics for Protein Domain Prediction. PLoS Comput Biol. 2015 Nov 17;11(11):e1004509. doi: 10.1371/journal.pcbi.1004509. PMID: 26575353; PMCID: PMC4648515.

123. Okamura K, Feuk L, Marquès-Bonet T, Navarro A, Scherer SW. Frequent appearance of novel protein-coding sequences by frameshift translation. Genomics. 2006 Dec;88(6):690–697. doi: 10.1016/j.ygeno.2006.06.009.

124. Pan X et al (2006) A DNA integrity network in the yeast Saccharomyces cerevisiae. Cell 124(5):1069–1081

125. Parikh SB, Houghton C, Branden Van Oss S, Wacholder A, Carvunis AR. Origins, evolution, and physiological implications of de novo genes in yeast. Yeast. 2022 Aug 12. doi: 10.1002/yea.3810. Epub ahead of print. PMID: 35959631.

126. Pasek S, Risler JL, Brézellec P. Gene fusion/fission is a major contributor to evolution of multi-domain bacterial proteins. Bioinformatics. 2006 Jun 15;22(12):1418–23. doi: 10.1093/bioinformatics/btl135. Epub 2006 Apr 6. PMID: 16601004.

127. Patraquim P, Magny EG, Pueyo JI, Platero AI, Couso JP. Translation and natural selection of micropeptides from long non-canonical RNAs. Nat Commun. 2022 Oct 31;13(1):6515. doi: 10.1038/s41467-022-34094-y.

128. Parvathy ST, Udayasuriyan V, Bhadana V. Codon usage bias. Mol Biol Rep. 2022 Jan;49(1):539–565. doi: 10.1007/s11033-021-06749-4.

129. Pearson WR. An introduction to sequence similarity (“homology”) searching. Curr Protoc Bioinformatics. 2013 Jun;Chapter 3:Unit3.1. doi: 10.1002/0471250953.bi0301s42. PMID: 23749753; PMCID: PMC3820096.

130. Phansalkar R, Lapierre P, Mellone BG. Evolutionary insights into the role of the essential centromere protein CAL1 in Drosophila. Chromosome Res. 2012 Jul;20(5):493–504. doi: 10.1007/s10577-012-9299-7. PMID: 22820845.

131. Piehler AP, Wenzel JJ, Olstad OK, Haug KB, Kierulf P, Kaminski WE. The human ortholog of the rodent testis-specific ABC transporter Abca17 is a ubiquitously expressed pseudogene (ABCA17P) and shares a common 5’ end with ABCA3. BMC Mol Biol. 2006 Sep 12; 7:28. doi: 10.1186/1471-2199-7-28. PMID: 16968533; PMCID: PMC1579226.

132. Pink RC, Wicks K, Caley DP, Punch EK, Jacobs L, Carter DR. Pseudogenes: pseudo functional or key regulators in health and disease? RNA. 2011 May;17(5):792–8. doi: 10.1261/rna.2658311. Epub 2011 Mar 11. PMID: 21398401; PMCID: PMC3078729.

133. Ponce R, Hartl DL. The evolution of the novel Sdic gene cluster in Drosophila melanogaster. Gene. 2006 Jul 19;376(2):174–83. doi: 10.1016/j.gene.2006.02.011.

134. Poretti M, Praz CR, Sotiropoulos AG, Wicker T. A survey of lineage-specific genes in *Triticeae* reveals de novo gene evolution from genomic raw material. Plant Direct. 2023 Mar 16;7(3):e484. doi: 10.1002/pld3.484.

135. Prade VM, Gundlach H, Twardziok S, Chapman B, Tan C, Langridge P, Schulman AH, Stein N, Waugh R, Zhang G, Platzer M, Li C, Spannagl M, Mayer KFX. The pseudogenes of barley. Plant J. 2018 Feb;93(3):502–514. doi: 10.1111/tpj.13794.

136. Reinhardt JA, Wanjiru BM, Brant AT, Saelao P, Begun DJ, Jones CD. De novo ORFs in Drosophila are important to organismal fitness and evolved rapidly from previously non-coding sequences. PLoS Genet. 2013; 9(10):e1003860. doi: 10.1371/journal.pgen.1003860.

137. Reva BA, Finkelstein AV, Skolnick J. What is the probability of a chance prediction of a protein structure with an rmsd of 6 A? Fold Des. 1998;3(2):141–7. doi: 10.1016/s1359-0278(98)00019-4.

138. Rojas-Duran MF, Gilbert WV. Alternative transcription start site selection leads to large differences in translation activity in yeast. RNA. 2012 Dec;18(12):2299–305. doi: 10.1261/rna.035865.112. Epub 2012 Oct 25. PMID: 23105001; PMCID: PMC3504680.

139. Rost B. Twilight zone of protein sequence alignments. Protein Eng. 1999 Feb;12(2):85–94. doi: 10.1093/protein/12.2.85. PMID: 10195279

140. Ruiz-Orera J, Messeguer X, Subirana JA, Alba MM (2014) Long non-coding RNAs as a source of new peptides. eLife 3, e03523

141. Samusik N, Krukovskaya L, Meln I, Shilov E, KozlovAP (2013) PBOV1 is a human de novo gene with tumor-specific expression that is associated with a positive clinical outcome of cancer. PLoS ONE 8, e56162

142. Savojardo C, Bruciaferri N, Tartari G, Martelli PL, Casadio R. DeepMito: accurate prediction of protein sub-mitochondrial localization using convolutional neural networks. Bioinformatics. 2020 Jan 1;36(1):56–64. doi: 10.1093/bioinformatics/btz512.

143. Schejter ED, Shilo BZ. Characterization of functional domains of p21 ras by use of chimeric genes. EMBO J. 1985 Feb;4(2):407–12. doi: 10.1002/j.1460-2075.1985.tb03643.x. PMID: 3926484; PMCID: PMC554200.

144. Schlötterer C (2015) Genes from scratch — the evolutionary fate of de novo genes. Trends Genet. 31, 215–2

145. Schmitz JF, Bornberg-Bauer E. Fact or fiction: updates on how protein-coding genes might emerge *de novo* from previously non-coding DNA. F1000Res. 2017 Jan 19;6:57. doi: 10.12688/f1000research.10079.1. PMID: 28163910; PMCID: PMC5247788.

146. Siepel A. Darwinian alchemy: Human genes from noncoding DNA. Genome Res. 2009 Oct;19(10):1693–5. doi: 10.1101/gr.098376.109.

147. Sievers F, Wilm A, Dineen D, Gibson TJ, Karplus K, Li W, Lopez R, McWilliam H, Remmert M, Söding J, Thompson JD, Higgins DG. Fast, scalable generation of high quality protein multiple sequence alignments using Clustal Omega. Mol Syst Biol. 2011; 7:539. doi: 10.1038/msb.2011.75.

148. Stoltzfus A. On the possibility of constructive neutral evolution. J Mol Evol. 1999 Aug;49(2):169–81. doi: 10.1007/pl00006540. PMID: 10441669.

149. Stothard P (2000) The Sequence Manipulation Suite: JavaScript programs for analyzing and formatting protein and DNA sequences. Biotechniques 28:1102–1104

150. Struhl K. The DNA-binding domains of the jun oncoprotein and the yeast GCN4 transcriptional activator protein are functionally homologous. Cell. 1987 Sep 11;50(6):841–6. doi: 10.1016/0092-8674(87)90511-3. PMID: 3040261.

151. Sutherland JM, Siddall NA, Hime GR, McLaughlin EA. RNA binding proteins in spermatogenesis: an in depth focus on the Musashi family. Asian J Androl. 2015 Jul-Aug;17(4):529–36. doi: 10.4103/1008-682X.151397.

152. Svensson EI, Berger D. The Role of Mutation Bias in Adaptive Evolution. Trends Ecol Evol. 2019 May;34(5):422–434. doi: 10.1016/j.tree.2019.01.015. PMID: 31003616.

153. Tautz D. The discovery of de novo gene evolution. Perspect Biol Med. 2014 Winter;57(1):149–61. doi: 10.1353/pbm.2014.0006. PMID: 25345708.

154. Tautz D, Domazet-Lošo T. The evolutionary origin of orphan genes. Nat Rev Genet. 2011 Aug 31;12(10):692–702. doi: 10.1038/nrg3053. PMID: 21878963.

155. Templeton AR. Has human evolution stopped? Rambam Maimonides Med J. 2010 Jul 2;1(1):e0006. doi: 10.5041/RMMJ.10006. PMID: 23908778; PMCID: PMC3721656.

156. Troskie RL, Faulkner GJ, Cheetham SW. Processed pseudogenes: A substrate for evolutionary innovation: Retrotransposition contributes to genome evolution by propagating pseudogene sequences with rich regulatory potential throughout the genome. Bioessays. 2021 Nov;43(11):e2100186. doi: 10.1002/bies.202100186.

157. Turelli P, Playfoot C, Grun D, Raclot C, Pontis J, Coudray A, Thorball C, Duc J, Pankevich EV, Deplancke B, Busskamp V, Trono D. Primate-restricted KRAB zinc finger proteins and target retrotransposons control gene expression in human neurons. Sci Adv. 2020 Aug 28;6(35):eaba3200. doi: 10.1126/sciadv.aba3200.

158. Uhlén, et al F. Proteomics. Tissue-based map of the human proteome. Science. 2015 Jan 23;347(6220):1260419. doi: 10.1126/science.1260419. PMID: 25613900.

159. Vakirlis N, Carvunis AR, McLysaght A (2020) Synteny-based analyses indicate that sequence divergence is not the main source of orphan genes. eLife 9

160. Vakirlis N, Hebert AS, Opulente DA, Achaz G, Hittinger CT, Fischer G, Coon JJ, Lafontaine I. A Molecular Portrait of De Novo Genes in Yeasts. Mol Biol Evol. 2018 Mar 1;35(3):631–645. doi: 10.1093/molbev/msx315. PMID: 29220506;

161. Vakirlis N, Vance Z, Duggan KM, McLysaght A. De novo birth of functional microproteins in the human lineage. Cell Rep. 2022 Dec 20;41(12):111808. doi: 10.1016/j.celrep.2022.111808. PMID: 36543139.

162. Van Oss SB, Carvunis AR. De novo gene birth. PLoS Genet. 2019 May 23;15(5):e1008160. doi: 10.1371/journal.pgen.1008160. PMID: 31120894; PMCID:

163. Vinckenbosch N, Dupanloup I, Kaessmann H. Evolutionary fate of retroposed gene copies in the human genome. Proc Natl Acad Sci U S A. 2006 Feb 28;103(9):3220–5. doi: 10.1073/pnas.0511307103. Epub 2006 Feb 21.

164. Walsh I, Seno F, Tosatto SC, Trovato A. PASTA 2.0: an improved server for protein aggregation prediction. Nucleic Acids Res. 2014 Jul;42(Web Server issue):W301–7. doi: 10.1093/nar/gku399. Epub 2014 May 21. PMID: 24848016; PMCID: PMC4086119.

165. Wang W, Yu H, Long M. Duplication-degeneration as a mechanism of gene fission and the origin of new genes in Drosophila species. Nat Genet. 2004 May;36(5):523–7. doi: 10.1038/ng1338. Epub 2004 Apr 4. PMID: 15064762.

166. Waterhouse A, Bertoni M, Bienert S, Studer G, Tauriello G, Gumienny R, Heer FT, de Beer TAP, Rempfer C, Bordoli L, Lepore R, Schwede T. SWISS-MODEL: homology modelling of protein structures and complexes. Nucleic Acids Res. 2018 Jul 2;46(W1):W296–W303. doi: 10.1093/nar/gky427.

167. Weisman CM, Murray AW, Eddy SR (2020) Many, but not all, lineage-specific genes can be explained by homology detection failure. PLoS Biol 18(11):e3000862

168. Weisman, C.M. The Origins and Functions of De Novo Genes: Against All Odds?. J Mol Evol 90, 244–257 (2022) 10.1007/s00239-022-10055-3

169. Weiss MC, Preiner M, Xavier JC, Zimorski V, Martin WF. The last universal common ancestor between ancient Earth chemistry and the onset of genetics. PLoS Genet. 2018 Aug 16;14(8):e1007518. doi: 10.1371/journal.pgen.1007518. PMID: 30114187

170. Wilson BA, Foy SG, Neme R, Masel J. Young Genes are Highly Disordered as Predicted by the Preadaptation Hypothesis of *De Novo* Gene Birth. Nat Ecol Evol. 2017 Jun;1(6):0146–146. doi: 10.1038/s41559-017-0146.

171. Williams SG, Lovell SC. The effect of sequence evolution on protein structural divergence. Mol Biol Evol. 2009 May;26(5):1055–65. doi: 10.1093/molbev/msp020.

172. Wissler L, Gadau J, Simola DF, Helmkampf M, Bornberg-Bauer E. Mechanisms and dynamics of orphan gene emergence in insect genomes. Genome Biol Evol. 2013;5(2):439–455. doi:10.1093/gbe/evt009

173. Wu DD, Irwin DM, Zhang YP. De novo origin of human protein-coding genes. PLoS Genet. 2011 Nov;7(11):e1002379. doi: 10.1371/journal.pgen.1002379

174. Yamashita Y, Nakamura N, Omiya K, Nishikawa J, Kawahara H, Obata H. Identification of an antifreeze lipoprotein from Moraxella sp. of Antarctic origin. Biosci Biotechnol Biochem. 2002 Feb;66(2):239–47. doi: 10.1271/bbb.66.239. PMID: 11999394.

175. Yan J, Kurgan L. DRNApred, fast sequence-based method that accurately predicts and discriminates DNA- and RNA-binding residues. Nucleic Acids Res. 2017 Jun 2;45(10):e84. doi: 10.1093/nar/gkx059. PMID: 28132027; PMCID: PMC5449545.

176. Yang W, Ng P, Zhao M, Wong TK, Yiu SM, Lau YL. Promoter-sharing by different genes in human genome--CPNE1 and RBM12 gene pair as an example. BMC Genomics. 2008 Oct 3;9:456. doi: 10.1186/1471-2164-9-456. PMID: 18831769; PMCID: PMC2568002.

177. Yang N, Zhao B, Chen Y, D’Alessandro E, Chen C, Ji T, Wu X, Song C. Distinct Retrotransposon Evolution Profile in the Genome of Rabbit (Oryctolagus cuniculus). Genome Biol Evol. 2021 Aug 3;13(8):evab168. doi: 10.1093/gbe/evab168.

178. Zhang Q, Backström N. Assembly errors cause false tandem duplicate regions in the chicken (Gallus gallus) genome sequence. Chromosoma. 2014 Mar;123(1-2):165–8. doi: 10.1007/s00412-013-0443-8. Epub 2013 Nov 10. PMID: 24213641.

179. Zhang J, Ghadermarzi S, Katuwawala A, Kurgan L. DNAgenie: accurate prediction of DNA-type-specific binding residues in protein sequences. Brief Bioinform. 2021 Nov 5;22(6):bbab336. doi: 10.1093/bib/bbab336. PMID: 34415020.

180. Zhang Z, Harrison PM, Liu Y, Gerstein M. Millions of years of evolution preserved: a comprehensive catalog of the processed pseudogenes in the human genome. Genome Res. 2003 Dec;13(12):2541–58. doi: 10.1101/gr.1429003.

181. Zhang Y, Hou L. Alternate Roles of Sox Transcription Factors beyond Transcription Initiation. Int J Mol Sci. 2021 May 31;22(11):5949. doi: 10.3390/ijms22115949.

182. Zhang L, Ren Y, Yang T, Li G, Chen J, Gschwend AR, Yu Y, Hou G, Zi J, Zhou R, Wen B, Zhang J, Chougule K, Wang M, Copetti D, Peng Z, Zhang C, Zhang Y, Ouyang Y, Wing RA, Liu S, Long M. Rapid evolution of protein diversity by de novo origination in Oryza. Nat Ecol Evol. 2019 Apr;3(4):679–690. doi: 10.1038/s41559-019-0822-5.

183. Zhang Y, Skolnick J. TM-align: a protein structure alignment algorithm based on the TM-score. Nucleic Acids Res. 2005 Apr 22;33(7):2302–9. doi: 10.1093/nar/gki524.

184. Zhang JY, Zhou Q. On the Regulatory Evolution of New Genes Throughout Their Life History. Mol Biol Evol. 2019 Jan 1;36(1):15–27. doi: 10.1093/molbev/msy206.

185. Zheng M, Chen X, Cui Y, Li W, Dai H, Yue Q, Zhang H, Zheng Y, Guo X, Zhu H. TULP2, a New RNA-Binding Protein, Is Required for Mouse Spermatid Differentiation and Male Fertility. Front Cell Dev Biol. 2021 Feb 18; 9:623738. doi: 10.3389/fcell.2021.623738.

186. Zhuang X, Cheng CC. Propagation of a De Novo Gene under Natural Selection: Antifreeze Glycoprotein Genes and Their Evolutionary History in Codfishes. Genes (Basel). 2021 Nov 9;12(11):1777. doi: 10.3390/genes12111777

187. Zhuang X et al (2019) Molecular mechanism and history of non-sense to sense evolution of antifreeze glycoprotein gene in northern gadids. Proc Natl Acad Sci 116(10):4400–4405

188. Zhou BB, Elledge SJ. The DNA damage response: putting checkpoints in perspective. Nature. 2000 Nov 23;408(6811):433–9. doi: 10.1038/35044005. PMID: 11100718.

