## Supplementary Figures for "Four classic “de novo” genes all have plausible homologs and likely evolved from retro-duplicated or pseudogenic sequences"

**Molecular Genetics and Genomics**

```
     M   F   P   R   P   V   L   N   S   R   A   Q   A   I   L
FLJ ATG-TTT-CCT-AGG-CCT-GTC-TTG-AAC-TCC-CGG-GCT-CAA-GCG-ATT-CTC    45
     M   L   P   R   L   V   L   N   S   L   P   Q   V   I   L
ZNF ATG-TTG-CCC-AGG-CTG-GTC-TTG-AAC-TCC-TTC-CCT-CAA-GTG-ATT-CTT    90

    *** **  **  *** *   *** *** *** ***      ** *** * * *** **


     L   P   Q   P   P   N   M   L   D   H   R       Q   W   P
FLJ CTG-CCT-CAG-CCT-CCC-AAC-ATG-CTG-GAT-CAC-AGG-----CAG-TGG-CCA    87
     L   P   N   P   P   K   V   L           R       Q   G   L
ZNF CTG-CCT-AAT-CCT-CCC-AAA-GTT-CTT---------AGA-TTA-CAG-GGA-CTA   129

    *** ***  *  *** *** **   *  **          **      ***  *  * *
```

**Suppl Fig 1:** The alignment of the N-termini of FLJ33706 and ZNF678.1 in humans, corresponding to the start site of the former which is located downstream in the latter. The codons shown in bold are those that are identical in both sequences, in total 13 out of 30. The 90 nt sequences shown are 68% identical with asterisks denoting each site. The translated sequences are also displayed. Underlined is the TGG codon which is a stop site, TAG, in other primates and so represents a premature termination.

```
FLJ AGTGCCAG-ACTGCA-GCT-GCCACATGAAGAGAGgt...agAGAGGG---TCTC-CT

      M  P  G

ZNF A-TGCCAGGACACCCCGAAAGCC----GGAAAACGgt...agAGAC-GCATTCTCACT    45

    * ****** **  *  *   ***    * * *  *       *** **   **** **
```

**Suppl Fig 2:** The alignment of part of the 5'UTR of FLJ33706 and the first 15 codons of the N-terminus of ZNF678, isoform 1, in humans showing 57% identity. The Kozak sequence of FLJ33706 was created by 2 indels that deleted 4 nucleotides. An insertion of guanine (underlined) within the original start site led to translation being initiated at a methionine codon further downstream. The proline (CCA) and most of the glycine (GGA) codon of ZNF678.1, shown in blue, is still found in FLJ33706. In bold are the splices sites found in the sequences that correspond with the positions of their respective introns. Shaded in pink is the part of the Kozak sequence immediately upstream of the start codon in FLJ33706.

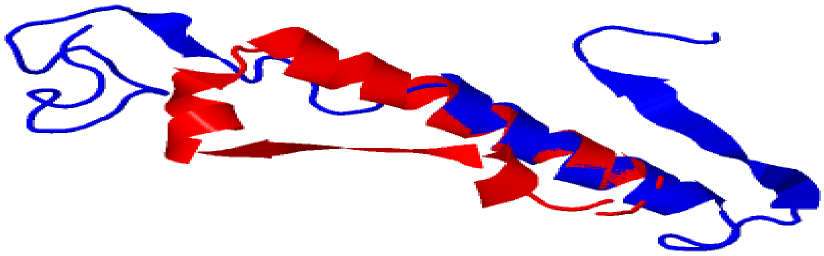

**Suppl Fig 3:** The predicted structural alignment of FLJ33706 (blue) and the corresponding sequence of ZNF678 (red) for the N-terminal region of the latter that contains a transcriptionally repressive KRAB domain. When superimposed, the peptides are highly similar in their atomic positions at least for the alpha helical structure but then tend to diverge in other places. The overall RMSD is calculated as 2.93

```
GDRD   --GATC--TGTTTGCA---CTTAAGTC-TGTTCTAGTTCACAGCTCC-TGA-ATAACTAT  50

PMAX   TTGGTCACTCT--GCAAGTCTTACAACGTGAGC-AGACAACAACAAAATTACATGTTTAT  57

         * **  * *  ***   ****   * **  * **   *** *    * * **   ***

GDRD   T-G-A-------------------------------------------------------  53

PMAX   TTGGAATAATCACAAAAAACAAAAAACTTACCAACTTAGTGGAATAATCTTAGGAAATAT 117

       * * *

GDRD   -----AATTTCTGAAATATATTTTTTT------TGT--CAAA                    82

PMAX   CTATTAATTTGCGCGCAATCCGTGCAAAACAAGTGAAGCAAC                   159

            *****  *    **   *          **   ***
```

**Suppl Fig 4:** The alignment of the 5'UTRs of Goddard and Protein MAX in D. melanogaster. There is a huge bulk deletion inferred in the former. Upstream of this deletion, there exists some similarity averaging about 50%, It is possible that enhancers, containing TF binding sites, are present. There is also a TATA box yet further upstream that could indicate the existence of an independent promoter

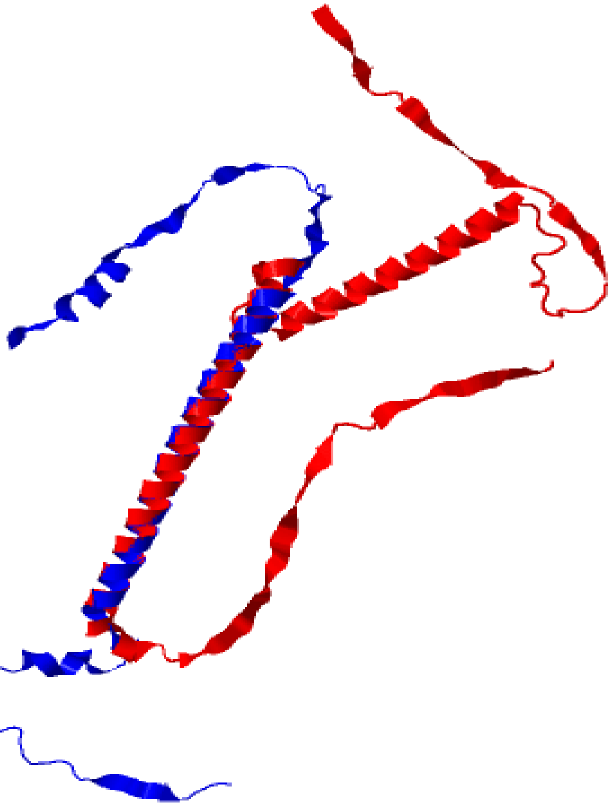

**Suppl Fig 5:** The predicted structural alignment of Protein MAX (red) and Goddard (blue) in D. melanogaster. These represent the full-length peptides. Goddard closely aligns with the Helix-Loop-Helix protein, MAX, in the main alpha helix. Both proteins are in the same fold except for where the model predicts a second helix the existence of which in Goddard is uncertain and may have been lost by premature truncation of the peptide sequence. The RMSD here is calculated as just 2.37 angstroms

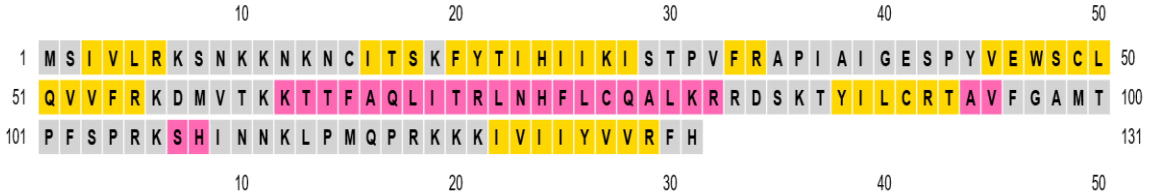


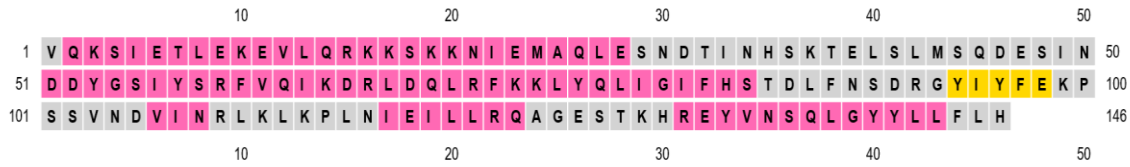


**Suppl Fig 6:** The predicted secondary protein structure of BSC4 in S. cerevisiae, above, and the corresponding sequence in Vps38. below, of the same yeast species. The latter consists mostly alpha helix (pink), but it does contain large sections of loop (grey) and some beta strand (yellow), the dominant structure of the former. The inferred transition to beta sheet would likely have been the result of a deformation which often can occur in protein evolution between alpha helix to beta sheet.

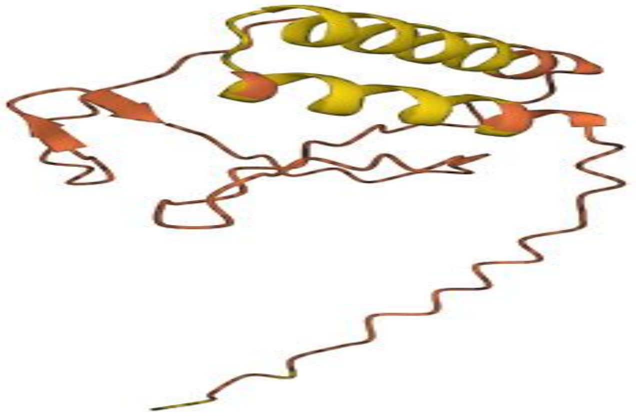

**Suppl Fig 7:** The predicted secondary protein structure of BSC4 in S. cerevisiae according to Alphafold. It reveals a much more stable, than rudimentary, conformation containing possibly two alpha helices.

| Protein type | Other | Signal peptide | Mitochondrial transfer peptide |
|---|---|---|---|
| Likelihood | 0.9766 | 0.0002 | 0.0232 |

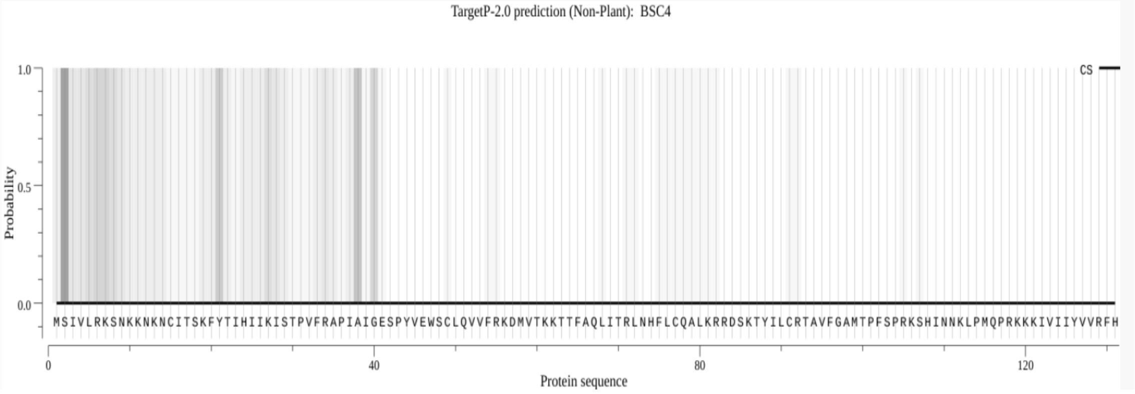


**Suppl Fig 8:** This plot by Target-2.0 indicates the detection of a weak, but still noticeable, signal for a mitochondrial transfer peptide at the N-terminus of BSC4. Using another tool, DeepMito, the predicted score is given as 0.18 compared to 0.43 for Vps38, that may also be localized to the vacuole

```
APOA2  CCACCACTACCCCCGCCCC-CACCCC-C-GA-CCAGACAG-TATAAA-------GAGTT   48

AFGP2  CTTGAA-TAAACCATCTATTCAACCCTCTGATCCA-ACTGGT-TAAGTGGCCTTGATTT   56

       *     * **   **   *     ****** * ** *** ** * * ***        ** **


APOA2  TACTCACGCTCCAGTCATTCAGTCTCCTGAGGACCCACTCCACCAGACCAA--CA-CC   105

AFGP2  --C-CAC-TTCAAG--A---A-T------A--A-CTACT--AC-ACTCCAAAGCAACC   98

         * ***   ** **   * * * *      *  * * ***  ** *  ****  ** **
```

**Suppl Fig 9:** The alignment of the entire 5’ UTRs of APOA-II in G. morhua and AFGP2 in the same cod species. It is fragmented in places but also highly similar in others, approximately 50% identical overall. Notably, the apolipoprotein 5’ UTR sequence has a much higher GC content (58%) than that of the afgp (40%). This may be explainable in terms of an AT mutational bias or because of natural selection.

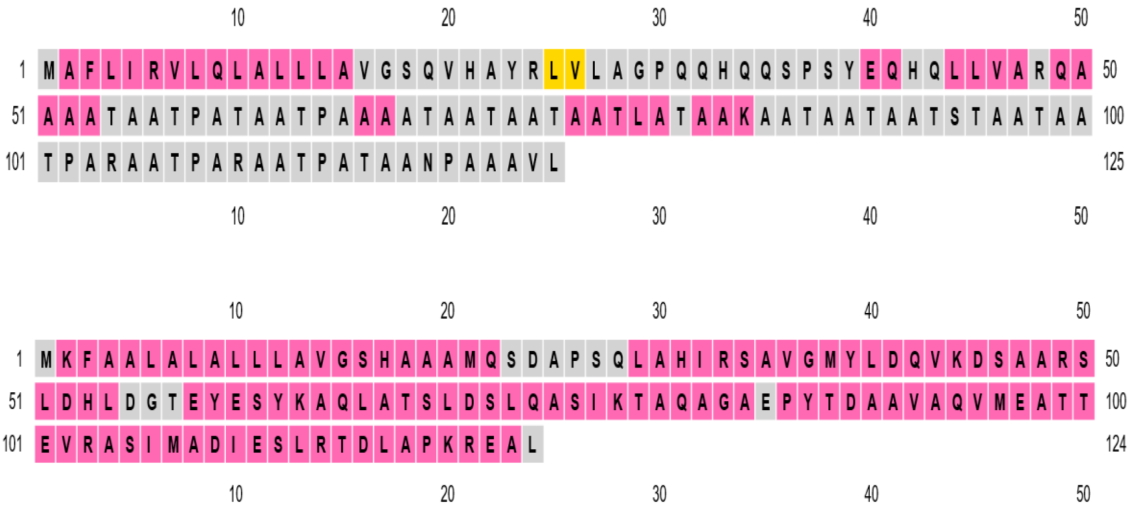


**Suppl Fig 10:** The predicted secondary protein structure of AFGP2 in G. morhua, above, and the corresponding sequence of APOA-II, below, in the same cod species. The latter is much more consistent, almost completed composed of alpha helices (shown in pink), whereas the former is more fragmented and variegated overall, and consists mainly of loop/coil (shown in grey) as well as a small amount of beta sheet (in yellow). Despite this, the afgp is not more intrinsically disordered of the two

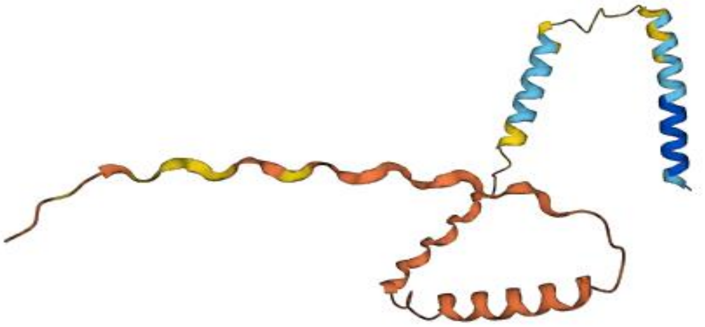

**Suppl Fig 11:** The predicted structure of AFGP2 in Alphafold revealing up to four short alpha helices

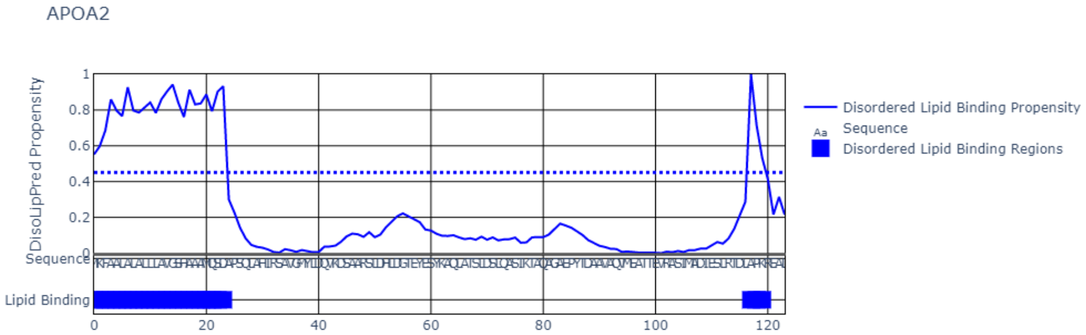


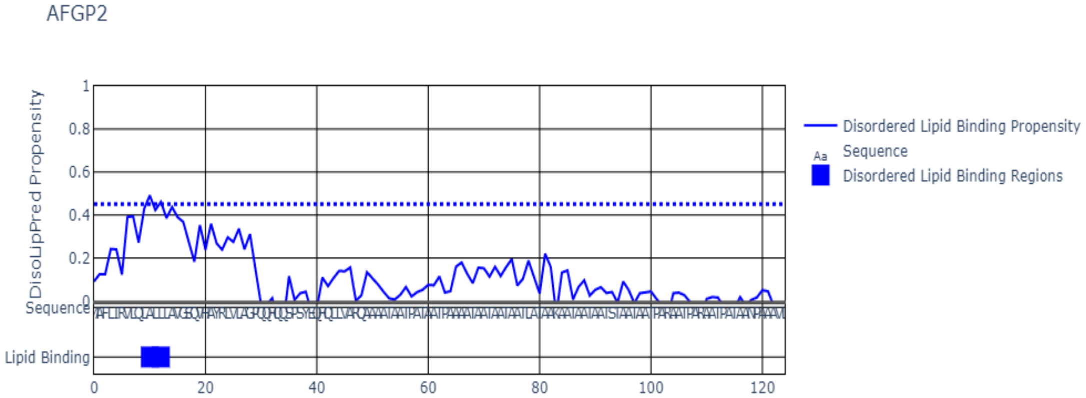


**Suppl Fig 12:** The predicted disordered lipid-binding propensity of APOA2 (above) and AFGP2 (below) (in G. morhua) for their aligned sequences determined using DisLipoPred. The latter appears to have retained a vestigial, albeit attenuated, capability inherited from its putative precursor. It allows it to bind with lipids at its N-terminal signal peptide. Previous research has indicated that the SP can itself interact with the hydrocarbon region of lipids. Lipoproteins, generally, also have anti-freeze potential.
